# Tumor Treating Fields Suppression of Ciliogenesis Enhances Temozolomide Toxicity

**DOI:** 10.1101/2021.12.02.470969

**Authors:** Ping Shi, Jia Tian, Brittany S. Ulm, Julianne C. Mallinger, Habibeh Khoshbouei, Loic P. Deleyrolle, Matthew R. Sarkisian

## Abstract

Tumor Treating Fields (TTFields) are low intensity, alternating intermediate frequency (200kHz) electrical fields that extend survival of glioblastoma patients receiving maintenance temozolomide (TMZ) chemotherapy. How TTFields exert efficacy on cancer over normal cells, or interact with TMZ is unclear. Primary cilia are microtubule-based organelles triggered by extracellular ligands, mechanical and electrical field stimulation, and are capable of promoting cancer growth and TMZ chemoresistance. We found in both low and high grade patient glioma cell lines that TTFields ablated cilia within 24 hours. Halting TTFields treatment led to recovered frequencies of elongated cilia. Cilia on normal primary astrocytes, neurons, and multiciliated/ependymal cells were less affected by TTFields. The TTFields-mediated loss of glioma cilia was partially rescued by chloroquine pretreatment, suggesting the effect is in part due to autophagy activation. We also observed death of ciliated cells during TTFields by live imaging. Notably, TMZ-induced stimulation of ciliogenesis in both adherent cells and gliomaspheres was blocked by TTFields. Moreover, the inhibitory effects of TTFields and TMZ on tumor cell recurrence correlated with the relative timing of TMZ exposure to TTFields and ARL13B^+^ cilia. Finally, TTFields disrupted cilia in patient tumors treated ex vivo. Our findings suggest TTFields efficacy may depend on the degree of tumor ciliogenesis and relative timing of TMZ treatment.

## Introduction

High grade gliomas in adult, such as glioblastoma (GBM), usually have dismal prognoses due to the resistance and recurrence following all standard of care treatments. These treatments include a combination of surgical resection (if possible), irradiation, and TMZ chemotherapy, the combination of which extends survival only a few months (1, 2), indicating novel treatments are urgently needed. One of the latest Food and Drug Administration-approved treatments for GBM patients is Tumor Treating Fields (TTFields) (Optune®), a device/electrode set that patients wear during their treatment that delivers low intensity (1-3V/cm), alternating intermediate frequency (200kHz) electric fields across the head. Combining maintenance treatment with TMZ, TTFields significantly increases overall survival several months beyond TMZ alone (3, 4). Thus TTFields are considered a new standard of care option (5). However, our understanding of how TTFields differentially targets gliomas over normal cells, interacts or enhances current therapies, or whether tumor cell characteristics predict sensitivity to TTFields remain unanswered questions.

The anti-tumor effects of TTFields do not occur via a single mechanism of action but rather a variety of cellular and molecular alterations (for review see: (6)). For example, TTFields disrupt the microtubular organization of mitotic spindle affecting normal cytokinesis and mitosis (7–9), as well as suppressing cell migration and invasion (10, 11). TTFields inhibit DNA damage repair and induce replication stress (12, 13). TTFields can induce autophagy (14) and promote immunogenic cell death (15). TTFields also change the cell plasma membrane permeability to a greater extent in tumor cells compared to primary dermal fibroblasts (16). Such membrane changes may be linked to calcium channel activation and rapid calcium influx that occurs during TTFields (17). Altogether these factors may cumulatively result in reduced proliferative and invasive capacity of glioma cells and enhanced sensitization to current therapies.

Many pathways impacted by TTFields are regulated by or involve signaling linked to the primary cilium (for review see: (18–20)). Primary cilia are non-motile, microtubule-based organelles extending from the mother centriole of the basal body. Cilia must be disassembled so centrioles within the basal body can duplicate, segregate and be re-purposed for mitosis. Cilia are ensheathed by plasma membrane distinct from membrane of the cell body (21–24), and generally depend on intraflagellar transport machinery for their outgrowth which mobilizes cargo anterogradely to the ciliary tip and retrogradely back to the cell body (25). At any given time, cilia are present on up to 30% or more of high-grade glioma cells (26). Nothing is known about how electrical field stimulation impact primary cilia on glioma cells. In human adipose-derived stem cells, brief exposures (4 hours/day) to low intensity (1V/cm), low frequency (1Hz) non-alternating electric fields were reported to induce osteogenesis via primary cilia (27). Electrical field-induced osteogenic responses were absent when ciliogenesis was inhibited using siRNA targeting an essential ciliogenesis gene intraflagellar transport 88 (*IFT88*)(27). Exposure to 16 Hz pulsed electromagnetic fields protected ciliary morphology against cigarette-smoke induced damage in osteoprogenitor cells (28). Thus, whatever role(s) primary cilia serve on glioma cells, they may be sensitive to or stimulated by the much higher therapeutic frequencies used in TTFields therapy.

The cilia on glioma cells may play a role in resistance to TMZ (29, 30). For example, cilia depletion mediated by CRISPR/Cas9 depletion of *PCM1* or *KIF3a,* two critical ciliogenesis genes, sensitized GBM cells to TMZ (29). More recently, TMZ was shown to induce enhancer of zeste homologue 2 (EZH2) which targets the expression of ADP ribosylation factor 13b (ARL13B), a regulatory GTPase highly concentrated in glioma cilia (26, 31), as an adaptive mechanism that promotes chemoresistance (30). Knockdown of ARL13b/cilia using shRNA in patient derived xenografts in vivo, not only slowed tumor growth, but increased sensitivity to TMZ in vivo. Thus, if TTFields affects ARL13B or ARL13B^+^ cilia, the sensitivity of glioma cells to TMZ could be enhanced. The goals of this study were to characterize whether and how TTFields at the clinical frequency (200kHz) affect glioma ciliogenesis compared to normal neural cell types in vitro. We also examined how TMZ alone versus TMZ plus TTFields affects ciliogenesis and proliferation on both ARL13b^+^ (ciliated) and ARL13B^-^ (non-ciliated) glioma cell lines characterized previously (32). Finally we explored whether TTFields affect ARL13B+ cilia in the patient tumor microenvironment.

## Results

### 1. Effects of different durations of TTFields exposure on low and high grade patient glioma cell primary ciliogenesis

We exposed two patient-derived glioma cell lines, L0 (a grade IV glioblastoma) and S7 (a grade II glioma) that grow primary cilia (26, 29, 31–33) to TTFields. We used Novocure’s Inovitro™ system to deliver low-intensity (1-4V/cm), 200kHz alternating electric fields to cultured cells which presumably mimic the type of fields delivered by the Optune® device in patients, similar to recent studies (16, 34). Generally, glioma cells were grown adherent (in serum) on coverslips or as free-floating spheres (without serum) for 3 days in vitro (DIV), then performed a single exposure to TTFields for up to 1 day or 3 continuous days at which point we analyzed cells immediately (‘acute’)(**Fig. 1A**). For repeated TTFields exposures we dissociated cells after 3 days of continuous TTFields and repeated the cycle 2 more times (**Fig. 1A**). For recovery, after the last day of single or repeated TTFields exposures, we dissociated spheres and cultured cells adherently (with serum) on coverslips and examined cells after 4-5 days.

**Figure 1.**
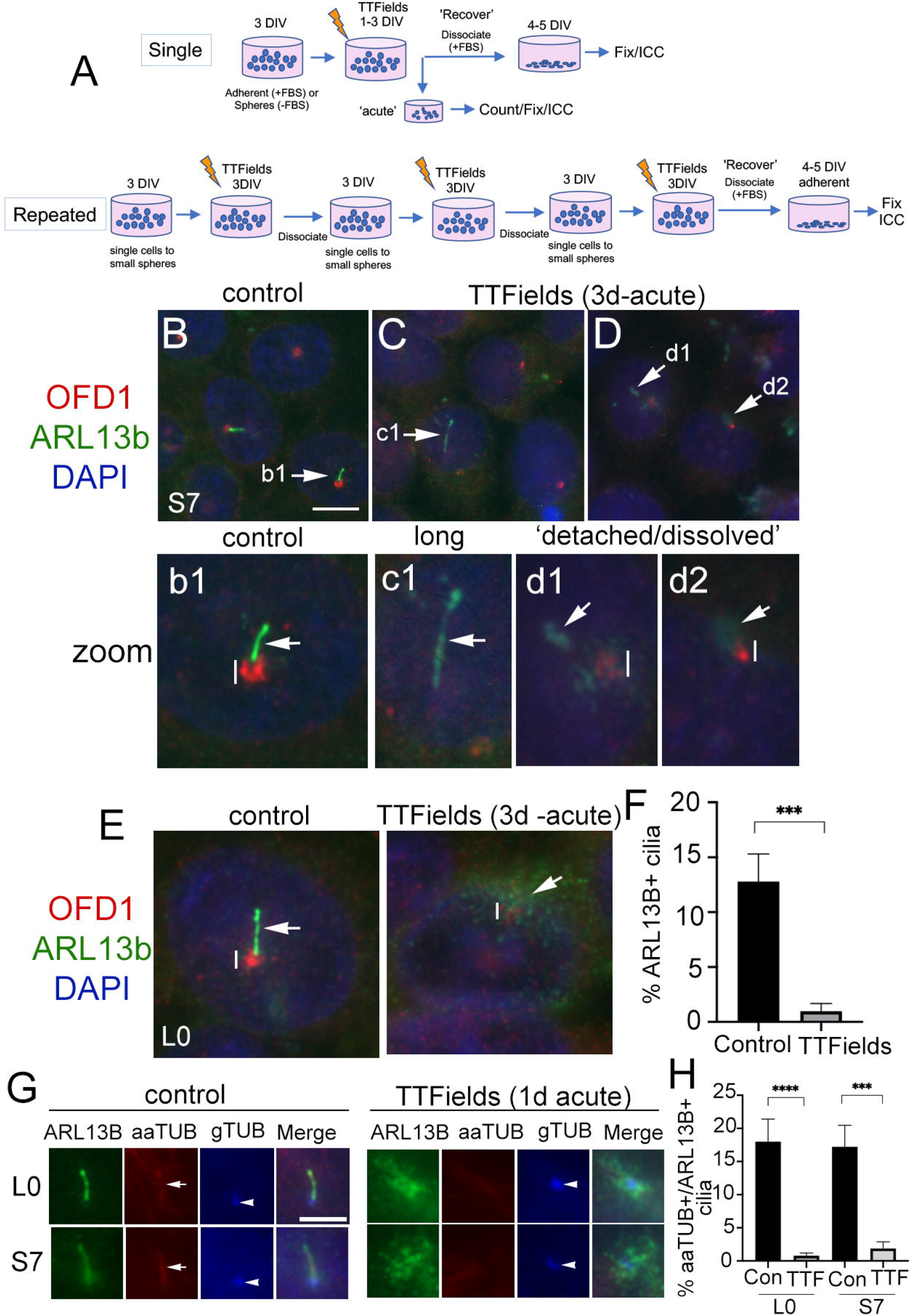
Primary cilia on patient-derived glioma cells are ablated by TTFields. **A**) General approach to treat cells with a single or repeated exposures to TTFields in vitro. Cells that were grown adherent on glass coverslips had media supplemented with 10% fetal bovine serum (FBS). **B-D**) S7 control cells stained for OFD1 (red), which clusters around basal bodies/centrioles, and ARL13B (green) which enriches in the primary cilium. Nuclei are labeled with DAPI (blue). In control (**B**), ARL13B^+^ cilia (arrow in b1 zoom) from OFD1^+^ puncta (vertical line). In S7 TTFields-treated cells (**C, D**) ARl13B^+^ cilia appeared elongated (e.g. **c1**), detached or separated away from the OFD1^+^ basal body (e.g. **d1**) or somewhat dissolved or cloudy in appearance at OFD1 clusters (e.g. **d2**). OFD1 clusters in TTFields-treated cells appeared smaller/less intense compared to control (compare vertical lines between b1 and d1/d2). **E**) Control or 3 days of TTFields-treated L0 cells stained similar to **B**. Control cells display clear OFD1+ basal body (vertical line) at the base of ARL13B^+^ cilia (arrow). TTFields-treated cells displayed less intensely-labeled OFD1^+^ basal body (vertical line) with surrounding dispersed/cloudy puncta of ARL13B (arrow). **F**) Percent of ARL13B^+^ cilia in L0 cells in control vs TTFields. **G**) L0 (upper row) and S7 (bottom row) cells stained for ARL13B (green), acetylated alpha tubulin (aaTUB, red), and gamma-tubulin (gTUB, blue). Control cells show ARL13B+ cilia colocalized with aaTUB (arrow) with gTUB^+^ basal bodies (arrowheads). TTFields-treated cells have clustered/dispersed ARL13B signal with no clear aaTUB^+^ axoneme associated with the gTUB^+^ basal body. **H**) Percent of aaTUB^+^/ARL13B^+^ cilia in control or TTFields-treated L0 and S7 cells. ***p<0.001, ****p<0.0001 (ANOVA). Scale bars (in μm) = 10 (**B**) and 5 (**G**).

After a single 3 days’ exposure to TTFields, we immunostained cells for ARL13B and orodigital facial syndrome 1 (OFD1), a protein that concentrates around the basal body (35–37). In control S7 cells, ARL13B+ cilia were readily identifiable extending from OFD1^+^ basal bodies (**Fig. 1B, b1**). After TTFields, the presence of ARL13B^+^ cilia were largely undetected. Cilia that remained were typically elongated (**Fig. 1C, c1**), or appeared detached from (**Fig. 1D, d1**) or dissolved (**Fig. 1D, d2**) around the basal bodies. Most TTFields-treated cilia displayed reduced intensity of OFD1 around the basal body compared to control (**Fig.1b1, d1, d2**). We observed similar phenomena in L0 cells (**Fig. 1E**). The appearance (**Fig. 1E**) and percent of ARL13B^+^ cilia (**Fig. 1F**) in L0 cells was significantly reduced. The effects of TTFields on glioma cilia can be seen within 24 hours post treatment. In both L0 and S7 cell lines there was a significant loss of ARL13B^+^ cilia after TTFields exposure (**Suppl. Fig. 1A-F**). To confirm TTFields are affecting the cilium and not just ARL13B localization along the ciliary membrane, we performed triple immunostaining to label a different component of cilia axoneme, acetylated-alpha tubulin (aaTUB) along with gamma-tubulin (gTUB) a microtubule component of the basal body/centriole, and ARL13B. In both L0 and S7 control cells, we found cilia that co-localized aaTUB^+^ and ARL13B^+^ extended from gTUB^+^ basal bodies (**Fig. 1G**). However after TTFields exposure, ARL13B puncta clustered around gTUB^+^ basal body/centrioles without obvious aaTUB^+^ axoneme extending from gTUB^+^ puncta (**Fig. 1G**). Quantification of cilia with both aaTUB^+^ and ARL13B^+^ cilia revealed significantly reduced frequencies after TTFields exposure in the two cell lines (**Fig. 1H**), indicating that TTFields disrupt the integrity of the entire organelle.

The above observations suggest TTFields effects on the cilia of glioma initiate within hours. Indeed, TTFields have been shown to disrupt glioma cell membrane permeability within the first hour of treatment (16). Thus, we examined the cilia axoneme and membrane 1 hour and 6 hours after TTFields using antibodies against aaTub, gTUB, ARL13B and inositol polyphosphate-5-phosphatase E (INPP5e). INPP5e localizes to the ciliary membrane where it interacts with ARL13B (22, 38). We found that after 6 hours cilia appeared longer than controls in both cell lines. The elongated ARL13B^+^ cilia displayed underlying colocalization with aaTUB^+^ suggesting that TTFields may stimulate a transient lengthening of the entire organelle within hours (**Suppl. Fig. 2A, C**). We also observed some anomalies in the ciliary distribution of ARL13B and INPP5e staining. ARL13B and INPP5e seemed to distribute evenly along the ciliary axoneme in control, but after 60 min and 6 hours exposure to TTFields, the staining pattern appear clustered or polarized toward the proximal and distal tips of the cilium (**Suppl. Fig. 2A, B, C**). In S7 cells we also observed unusual clusters of INPP5e surrounding the basal body after TTFields exposure (**Suppl. Fig. 2B**). Thus, in addition to a ciliary lengthening that precedes the loss of cilia, TTFields affect properties of the cilia membrane and surrounding base within hours of exposure in vitro.

The significant depletion of primary cilia by TTFields led us to ask if this effect was permanent. That is, would the frequency of ciliated glioma cells remain low if treatment is stopped and cells are allowed to recover? After single or repeated TTFields exposure, we plated cells in serum for 4-5 days, fixed, and immunostained for ARL13B and pericentriolar material 1 (PCM1), another protein that clusters around the basal body and centrioles in glioma cells (29). In S7 cells, ARL13B+ cilia were detectable but appeared longer than control (**Fig. 2A**). Quantification of lengths of ARL13B^+^ cilia demonstrated a significant increase in the TTFields exposed groups (**Fig. 2B, C**). A similar increase in length in L0 was observed after recovery from TTFields (**Fig. 2D**). However, in either S7 or L0 cells there was no change in cilia frequency after recovery from TTFields (**Fig. 2E-G**). These data indicate that frequencies of ciliated glioma cells are restored after TTFields but are affected in a way that leads to elongation.

**Figure 2.**
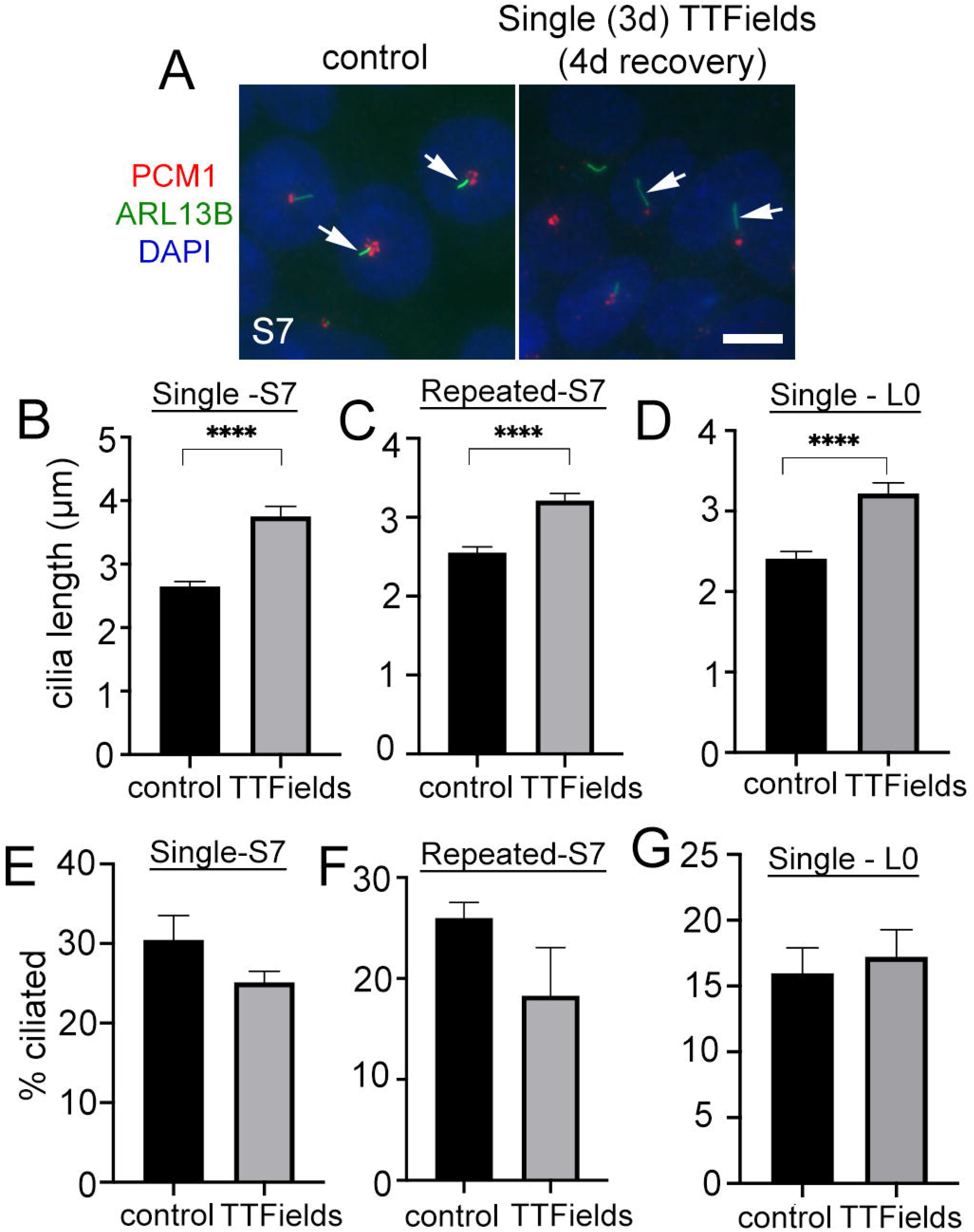
Similar frequencies of elongated glioma cilia after halting TTFields treatment. **A**) S7 cells with or without 3 days (3d) of TTFields treatment then dissociated onto coverslips for 4d. Fixed cells were immunostained for ARL13B (green) and PCM1 (red) with nuclei labeled with DAPI (blue). Control and TTFields-treated cells showed ARL13B^+^ cilia (arrows) extending from PCM1^+^ puncta which concentrates around basal bodies. Scale bar = 10μm. **B-D**) Mean lengths (μm) of S7 (**B, C**) and L0 (**D**) ARL13B^+^ cilia after 4 days recovery from a single or repeated exposure to TTFields. **E-G**) Percent of ciliated cells in S7 (**E, F**) and L0 (**G**) after 4 days recovery from a single or repeated exposure to TTFields. ****p<0.0001 (ANOVA).

### 2. TTFields do not have the same impact on normal mouse neural cilia

Considering the robust depletion of glioma cilia within 24 hours of exposure to TTFields, we next asked whether cilia of normal primary neural cell types are similarly affected by TTFields. To test this, we cultured dissociated mouse embryonic cortices on glass coverslips for 11 days in vitro (DIV). At 11DIV, we assigned coverslips as control or TTFields (24 hours or 3days exposure). One advantage of this type of culture is that we can examine the effects of TTFields on cells that differentiate into various subtypes including astrocytes and neurons, marked by glial fibrillary acidic protein (GFAP) and neuronal nuclei (NeuN) expression respectively.

We first examined astrocyte cilia through a combined immunostaining for GFAP, ARL13B and pericentrin (Pcnt, a protein concentrated around the cilia basal body) (**Fig. 3A, D**). After 24 hours of TTFields, we did not observe significant differences in the frequency (**Fig. 3B**) or length (**Fig. 3C**) of astrocyte cilia. However, after 3days of TTFields, there were significantly fewer ciliated GFAP^+^ cells (**Fig. 3E**), though the lengths of these cilia were comparable to control (**Fig. 3F**). These data suggest that, at least at acute timepoints after TTFields, astrocyte cilia appear more resistant to TTFields than glioma cells.

**Figure 3.**
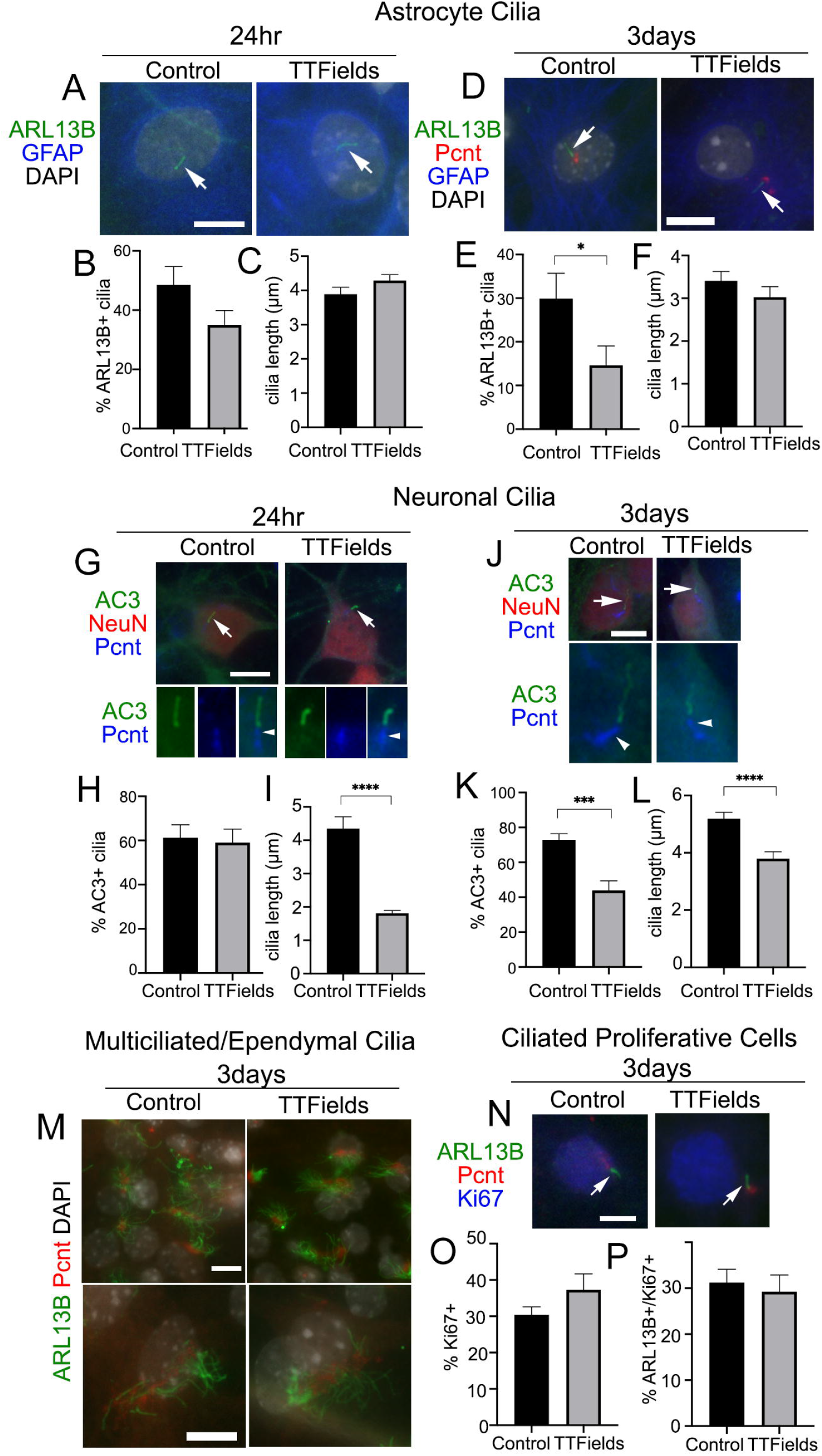
Effects of TTFields on normal mouse neural cell types in vitro. Mixed primary cultures from neonatal mouse cerebral cortex were dissociated and maintained for 11DIV and left untreated (Control) or exposed to 24 hour or 3 days continuous TTFields and fixed. **A**) Cells were stained for ARL13B (green) and GFAP (blue) after 24 hour TTFields. Nuclei were labeled with DAPI. Arrows point to ARL13B^+^ cilia in both groups at 24 hours. **B,C**) Percent of GFAP^+^ cells with ARL13B^+^ cilia (B), and mean length of ARL13B^+^ cilia (C) on GFAP^+^ cells after 24 hours. **D**) Cells were stained for ARL13B (green), pericentrin (Pcnt, red), and GFAP (blue) after 3 days TTFields. **E**) Percent of GFAP^+^ cells with ARL13B^+^ cilia (E), and mean length of ARL13B^+^ cilia (F) on GFAP^+^ cells after 3 days. **G**) Cells were stained for type 3 adenylyl cyclase (AC3) (green), NeuN (red), and Pcnt (blue) after 24 hours TTFields. Nuclei were labeled with DAPI. The arrows in upper panels point to cilia enlarged below each image showing Pcnt^+^ basal bodies for indicated cilia. **H, I**) Percent of NeuN^+^ cells with AC3^+^ cilia (H), and mean length of ARL13B^+^ cilia (I) on NeuN^+^ cells after 24 hours TTFields. **J**) Cells were stained for AC3 (green), NeuN (red), and Pcnt (blue) after 3 days TTFields. The arrows in upper panels point to cilia enlarged below each image which shows the Pcnt^+^ basal bodies for indicated cilia. **K**, **L**) Percent of NeuN^+^ cells with AC3^+^ cilia (**K**), and mean length of ARL13B^+^ cilia (**L**) on NeuN+ cells after 3 days TTFields. **M**) Cells were stained with ARL13B (green) and Pcnt (red) with nuclei labeled with DAPI (white). Lower magnification (upper panels) and enlarged (lower panels) examples of multiciliated cells in control (left panels) and 3 days of TTFields (right panels). Bars = 10μm **N**) Cells were stained for ARL13B (green), Pcnt (red) and Ki67 (blue) after 3 days TTFields. Images show examples of Ki67^+^ nuclei with ARl13B^+^ cilia (arrows) extending from Pcnt^+^ basal bodies. **O**) Percent of Ki67^+^ cells per field analyzed in each group. **P**) Percent Ki67^+^ cells with ARL13B^+^ cilia. * p<0.05, *** p<0.001, **** p<0.0001 (ANOVA).

Next we examined neuronal cilia by triple immunostaining for NeuN, type 3 adenylyl cyclase (AC3), an enzyme enriched in most neuronal cilia in the cortex (39–41), and Pcnt (42, 43) (**Fig. 3G, J**). After 24 hours of TTFields, we did not observe any significant changes on the frequency of neurons with AC3^+^ cilia (**Fig. 3H**), but the lengths of AC3^+^ cilia were significantly reduced (**Fig. 3I**). After 3 days of TTFields, both the frequency and length of AC3^+^ cilia on NeuN+ cells were significantly reduced compared to control (**Fig. 3K and L**, respectively). The extent of the reduced ciliary frequency was ~40% of control neurons, compared to about a 90% decrease in L0 cells after similar duration. Thus, neurons, especially after 24hr of TTFields, appear more resistant though not completely spared from the effects of TTFields.

The neural cultures also contained populations of multiciliated cell types (presumably cells that differentiated into ependymal cells) and proliferating cells. However the cells bearing tufts of cilia, detected by combination of ARL13B and Pcnt, appeared comparable after 24 hr (data not shown) and 3 days of continuous TTFields (**Fig. 3M**). A fraction of the cells in the culture were still Ki67^+^ suggesting they were still active in the cell cycle (**Fig. 3N**). However, amongst the Ki67^+^ cells (**Fig. 3O**) we did not observe significant change in the percentage of ciliated cells (**Fig. 3P**). Thus TTFields seems to spare the ability for multiciliated cells and ciliated cycling cells to form or maintain their cilia. Altogether these data suggest a differential sensitivity to TTFields between normal neural cell types and glioma cells.

### 3. TTFields-induction of autophagy and death of ciliated cells contribute to cilia depletion

What is the mechanism through which TTFields promotes rapid cilia loss in glioma cells? A number of factors and pathways promote cilia disassembly (44). Examples include calcium shock/influx (45), or autophagy activation (46), processes that have been shown to rapidly increase after TTFields onset (14, 17). We examined whether buffering extra/intracellular Ca^2+^ by pre-treating cells with 600mM EGTA or 1μM BAPTA increase cilia frequency during TTFields, however we did not observe any prevention of cilia loss (data not shown). We then examined if the autophagy pathway activation at cilia was involved, in part because the reduced OFD1 expression we observed around the basal bodies after TTFields, is a potential indicator of autophagy activation (37, 47). In addition, we observed microtubule-associated proteins 1A/1B light chain 3B (LC3B) and phospho-AMPK (pAMPK) recruitment to basal bodies after single and repeated TTFields application (**Suppl. Fig. 3**) consistent with reports that autophagy proteins are recruited to primary cilia (46, 48). Thus we pre-treated S7 and L0 cells 30 minutes before TTFields induction with vehicle or the autophagy inhibitor chloroquine (CQ) (20μM), fixed cells after 6 or 24 hours, and analyzed the frequency and length of cilia. A concentration of 20μM CQ was selected because it inhibited autophagy pathway activation in response to TTFields in U87 and other glioma cell lines (14). We found that S7 glioma cells pre-treated with CQ 30 minutes before TTFields led to a significant increase in the percent of ciliated cells after 24 hours compared to control (**Fig. 4A-D, E**). The TTFields-induced increase in cilia length in S7 cells was also reduced by CQ at 24hr (**Fig. 4G**). In L0 cells, we observed significantly more ciliated cells after 6 and 24 hours pretreatment with CQ and exposed to TTFields (**Fig. 4F**). In addition cilia length of CQ-treated L0 cells at 6 hours was significantly reduced compared to vehicle after 6 hours TTFields (**Fig. 4H**) suggesting autophagy activation may be underlying the observed elongation after TTFields. It is noteworthy that in both S7 and L0 cells, 6 hours of TTFields was sufficient to observe significant cilia elongation. Further, because CQ did not fully restore the frequency of cilia to control suggests that either CQ may not inhibit autophagy in all cells, or that the activation of autophagy may represent one of the factors resulting in TTFields-induced cilia depletion.

**Figure 4.**
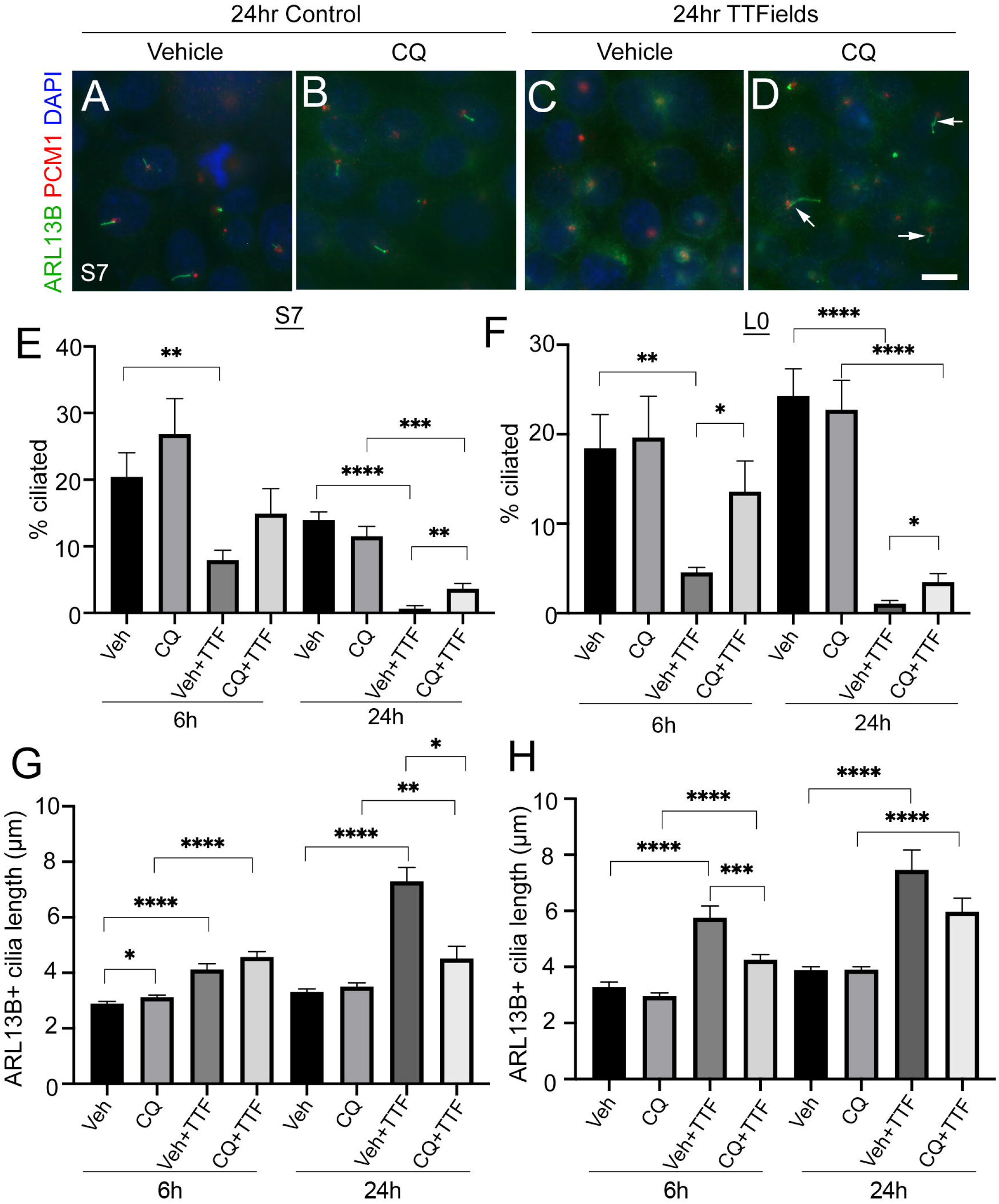
Chloroquine pretreatment partially prevents TTFields-induced loss of cilia. **A-D**) S7 cells pre-treated with vehicle (**A, C**) or 20μM chloroquine (CQ) (**B, D**) and left untreated (control) (**A,B**) or exposed to 24 hours TTFields (**C,D**). Cells were stained for ARL13B (green), PCM1 (red) with nuclei stained with DAPI (blue). Arrows in (D) point to cilia observed in cells treated for 24 hours of CQ+TTFields. Scale bar in D = 10μm. **E, F**) Percent of S7 (E) and L0 (F) cells with ARL13B^+^ cilia for the indicated treatment group after 6 hour (h) or 24 hours of TTFields (TTF). **G,H**) Mean lengths of ARL13B^+^ cilia in S7 (**G**) or L0 (**H**) cells for the indicated treatment group after 6 hours or 24 hours of TTFields (TTF). *p<0.05, **p<0.01, ***p<0.005, ****p<0.0001 (ANOVA).

To more directly examine how ciliated cells respond to TTFields, we transfected L0 and S7 cells with two cDNA constructs encoding ARL13B:GFP and OFD1:mCherry allowing us to track isolated cells displaying ARL13B:GFP^+^ cilia with OFD1:mCherry^+^ clusters around the basal body overnight (**Suppl. Fig. 4A**). We live imaged cells up to 24 hours after transfection using Novocure’s inovitro LIVE imaging system. Notably within several hours after TTFields onset we observed ciliated L0 cells that appeared to die (**Suppl. Fig. 4B, C**), with similar observations in S7 cells during TTFields (**Suppl. Fig. 4D**). These data indicate TTFields may have a direct impact on the survival and proliferation of ciliated glioma cells or cells that derive from ciliated glioma cells.

### 4. TMZ-induced ciliogenesis is blunted by TTFields

The survival benefit promoted by TTFields in patients occurs during TMZ maintenance therapy (3, 4). TMZ has recently been reported to increase the frequency and length of ARL13B+ cilia patient-derived glioblastoma cells (30). Thus we asked if the effects of TTFields on cilia would be similar in the presence of TMZ chemotherapy. The effects of TMZ on glioma ciliogenesis have not been extensively analyzed with respect to different cell lines, different concentrations and durations of exposure. Thus, we first examined how different durations and concentrations of TMZ, doses lower than those typically used to kill cells in in vitro assays, affect the frequency and length of ARL13B+ glioma cilia in our cell lines.

In L0 and S7 cells, 24 hours treatment with 10μM TMZ appeared to elongate primary cilia in both cell lines (**Fig. 5A**). In S7 cells, we observed a dose-dependent effect with 0.3, 3 and 10 μM TMZ sufficient to increase cilia length after 24 hours exposure (**Fig. 5B**). In L0 cells, we found that 10μM TMZ significantly increased the length of ARL13B^+^ cilia compared to lower TMZ concentrations and vehicle (**Fig. 5C**). After 3 days of exposure to TMZ (a duration chosen because we performed 3 days of continuous TTFields), we found that in S7 cells, 10μM and 50μM TMZ significantly reduced the length of cilia (**Fig. 5D**) but significantly increased the frequency of ciliated cells (**Fig. 5E**). However, 3 days of TMZ exposure in L0 cells did not affect cilia length after 10μM or 50μM (**Fig. 5F**), but significantly increased the frequency of ciliated L0 cells as in S7 cells (**Fig. 5G**). These results support results of recent studies (30) and suggest TMZ is capable of stimulating the elongation of ARL13B^+^ cilia, at least acutely, and increasing the frequency of ciliated glioma cells.

**Figure 5.**
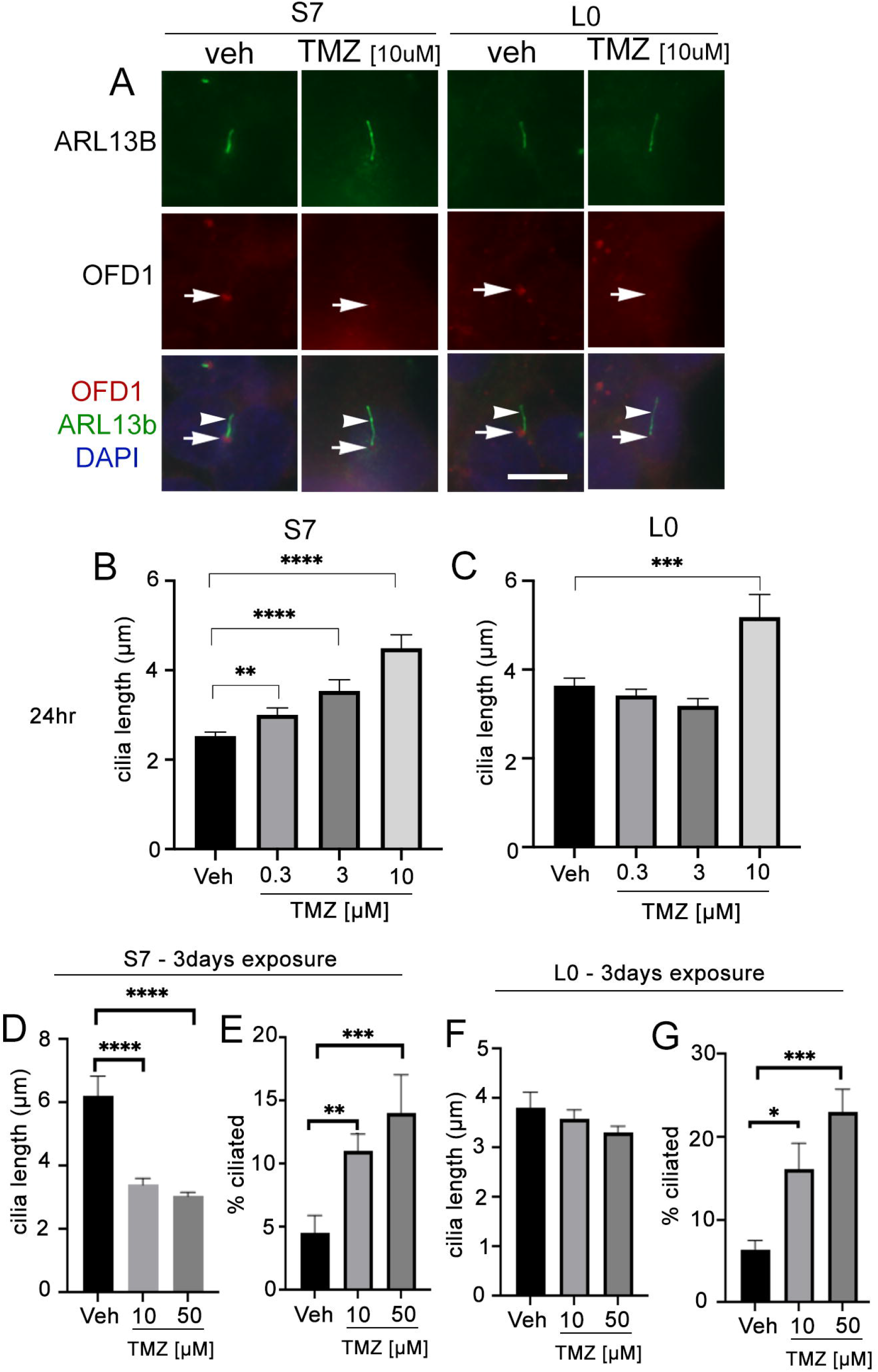
Temozolomide stimulates ciliogenesis in L0 and S7 cells. **A**) S7 (left two columns) and L0 (right two columns) were grown for 24 hours in vehicle or 10 μM TMZ. Cells were fixed and immunostained for OFD1 (red), ARL13B (green). Nuclei were labeled with DAPI. TMZ-treated cells appeared to show elongated ARL13B^+^ cilia (arrowheads) and OFD1 intensity at the base of the cilia (arrows) appeared to decrease compared to vehicle in both cell lines. Scale bar = 10μm. **B,C**) Mean lengths of cilia in S7 (**B**) and L0 (**C**) cells after a 24 hours exposure to the indicated concentration [μM] of TMZ. **D,F**) Mean lengths of ARL13B^+^ cilia in S7 (**D**) or L0 (**F**) cells after a 3 day exposure to the indicated concentration [μM] of TMZ. **E, G**) Percent of ARL13B^+^ cilia in S7 (**D**) or L0 (**F**) cells after a 3 day exposure to the indicated concentration [μM] of TMZ.

Since TMZ generally stimulates ciliogenesis, and TTFields inhibit it, we examined glioma cilia with a combination of these treatments in adherent cells and spheres (**Fig. 6**). First we pretreated adherent S7 and L0 cells with concentrations of TMZ that stimulated ciliogenesis about 30 minutes before a 3 days exposure to TTFields. We found that in both adherent S7 (**Fig. 6A-D**) and L0 (**Fig. 6E-H**) cells, ciliogenesis was not observed in the presence of TMZ plus TTFields (**Fig. 6D, H**). We also examined if TTFields had the same effect in gliomaspheres. We cultured S7 and L0 spheres for 3 days, and then treated them for 3 days with vehicle, 50μM TMZ, TTFields, or TTFields plus 50μM TMZ (added 30min before onset) (**Fig. 6I-L**). We then collected and fixed spheres, sectioned and immunostained them for ARL13B. For each sphere, we normalized the number of cilia to the area of the sphere. In S7 cells, we found that TMZ alone increased the frequency of cilia (**Fig 6M**), consistent with what we observed in adherent cells (**Fig. 5E**). As expected, TTFields significantly reduced the frequency of cilia in the spheres but TMZ-induced increase did not occur in the presence of TTFields (**Fig. 6M**). Interestingly we did not observe a change in the length of cilia across groups (**Fig. 6N**). Unlike S7 cells, TMZ alone did not increase the frequency of cilia in L0 spheres (**Fig. 6O**). However, like S7 cells, TTFields reduced the frequency of cilia in the L0 spheres which was not enhanced by the co-treatment of TMZ plus TTFields (**Fig. 6O**). Together, these results indicate that TTFields disrupt the pro-ciliogenic effects of TMZ.

**Figure 6.**
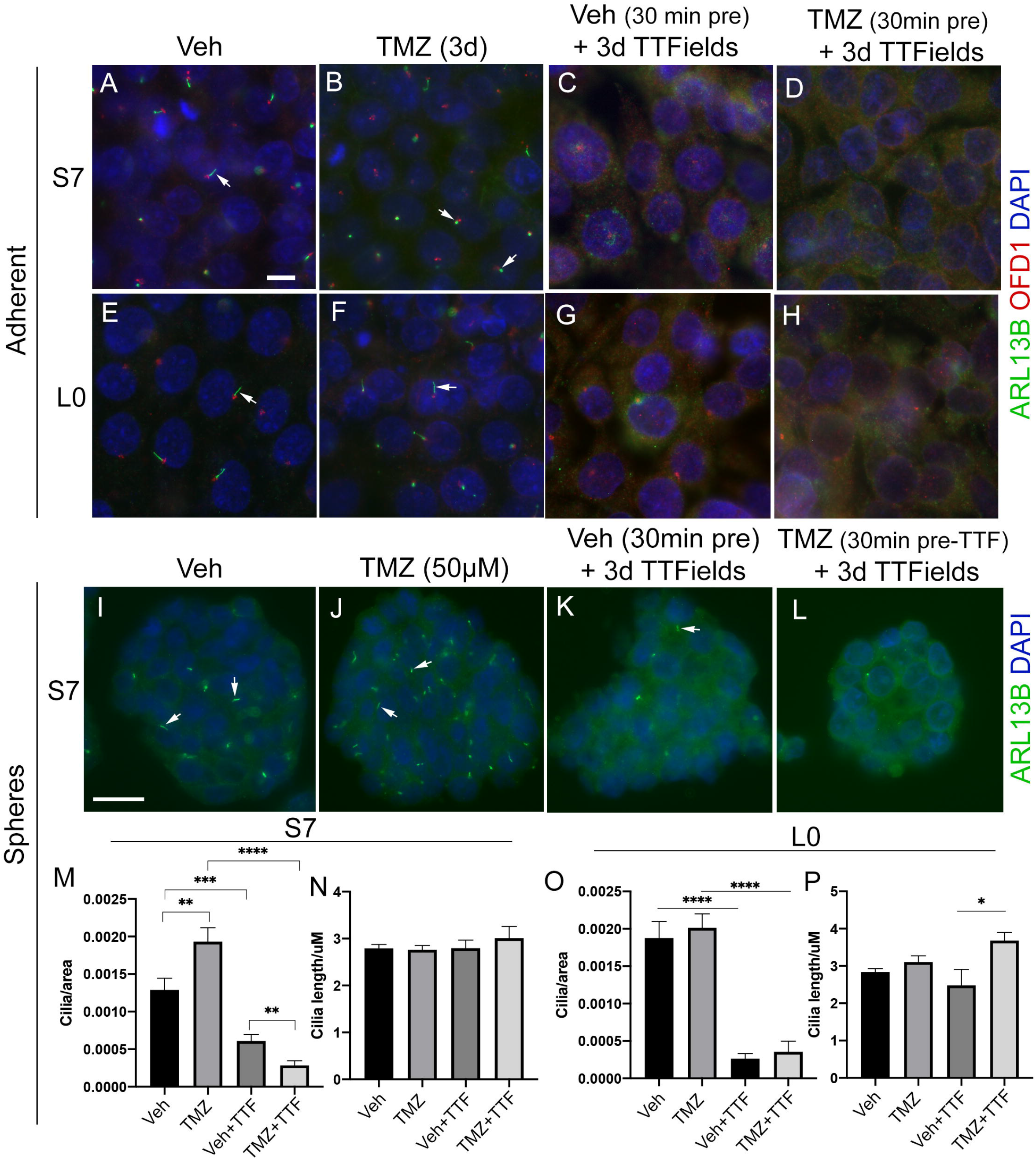
TTFields blocks the TMZ-mediated increase in ciliogenesis. **A-D**) Adherent S7 cells pre-treated with vehicle (**A, C**) or 10μM TMZ (**B, D**) and left untreated (control) (**A,B**) or exposed to 3 days (d) TTFields (**C,D**). **E-H**) Adherent L0 cells pre-treated with vehicle (**A, C**) or 20μM TMZ (**B, D**) and left untreated (control) (**A,B**) or exposed to 3 days TTFields. S7 and L0 cells were fixed and stained for ARL13B (green) and OFD1 (red). Nuclei are labeled with DAPI (blue). **I-L**) Spheres of S7 cells pre-treated with vehicle (**I, K**) or 50μM TMZ (**J, L**) and left untreated (control) (**I,J**) or exposed to 3 days TTFields (**K,L**). To examine cilia, spheres were fixed, sectioned and stained for ARL13B (green, arrows). **M, O**) The number of cilia/per area (μm^2^)of traced sections of S7 (M) or L0 (O) spheres for the indicated treatment. **N, P**) The mean length (μm) of cilia in S7 (N) or L0 (P) spheres for the indicated treatment. *p<0.05, **p<0.01, ***p<0.001, ****p<0.0001 (ANOVA). Scale bars (in μm) in A =10, I= 20.

### 5. The combined efficacy of TMZ and TTFields correlates with the timing of treatment and ARL13B^+^ cilia

Previously we found that deleting key ciliogenesis genes (e.g. *PCM1, KIF3A*) enhanced sensitivity of glioma cells to TMZ (29). Similarly, glioma cells expressing ARL13B shRNA, which depleted cilia, were more sensitized to TMZ in vitro and in vivo (30). Using our S7 glioma transgenic cell line depleted in ARL13B and cilia using CRISPR/Cas9 (32), we examined how these cells proliferated in response to TMZ, TTFields, or TMZ and TTFields co-treatment. After four days of growing S7 parental or ARL13B KO spheres, we treated them with vehicle, 50 or 100μM TMZ in the absence or with 3 days of TTFields (**Fig. 7A**). In parental S7 cells, proliferation was reduced in 100μM TMZ group (**Fig. 7B**). However in *ARL13B* KO cells, proliferation was significantly reduced at 50μM and 100μM TMZ groups (**Fig. 7B**), consistent with results of previous studies that ARL13B^+^ cilia are associated with resistance to TMZ. However, when we co-treated S7 parental or *ARL13B* KO cells with TMZ and TTFields, there was no added toxicity in either group (**Fig. 7C**). This suggests in cells with cilia ablation via treatment with TTFields or thru genetic means, the co-treatment of TMZ and TTFields may not lead to an acute additive toxicity.

**Figure 7.**
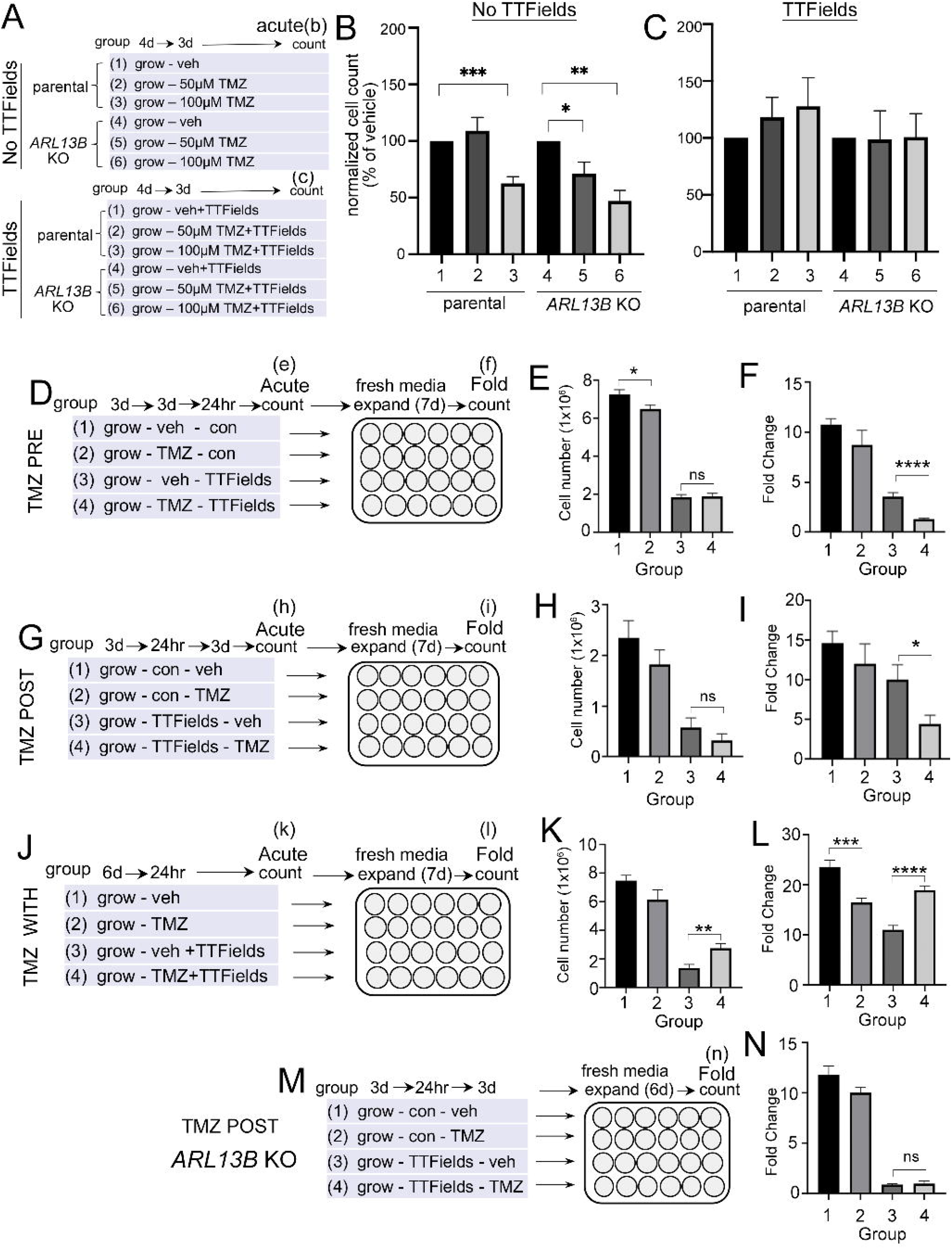
Combined TMZ and TTFields efficacy correlates with treatment sequence and ARL13B^+^ cilia. **A-C**) S7 parental or *ARL13B* KO (clone G12) cells were grown as spheres for 4 days and then treated with vehicle (veh), 50 or 100μM TMZ and exposed to an additional 3 days without (**B**) or with TTFields (**C**). Bar graphs show the normalized cell number (% of control) at the end of 7 days in the indicated group in A. **D)** TMZ ‘pre’ experiment: S7 parental cells were treated with 100μM TMZ for 3 days before a 24 hr treatment of TTFields or control (con). Cells were counted immediately after this treatment (Acute count) (**E**), and then pooled, and 2.5×10^4^ cells/well were expanded in fresh media for 7 days in 24 well plates (n=12 wells/group) and the fold expansion was calculated (Fold count) (**F**). **G-I**) TMZ ‘post’ experiment: TMZ (100μM) was given for 3 days after a 24hr TTFields treatment with S7 cells being counted acutely (H) or expanded in fresh media for 7 days (I). **J-L**) TMZ ‘with’ experiment: S7 cells were grown for 6 days and then simultaneously treated with TMZ (100μM) and TTFields for 24hrs, counted acutely (**K**) or examined for fold expansion after 7 days (**L**). **M**) Similar experiment as (G) but examining fold expansion of S7 *ARL13B* KO cells. ns =not significant, *p<0.05, **p<0.01, ***p<0.001, ****p<0.0001 (ANOVA)

Surprised that 3 days of TMZ and TTFields co-treatment in S7 parental cells did not show additive toxicity (**Fig. 7C**), we wondered if potential co-toxicity relates to treatment sequence or the treatment effect could be delayed. For example, would stimulating ciliogenesis with TMZ render more cells susceptible to TTFields, and/or would TTFields suppression of ciliogenesis sensitize more glioma cells to TMZ? To test this, we administered TMZ before (PRE) (**Fig. 7D-F**), after (POST) (**Fig. G-I**) or during (WITH) (**Fig. 7J-L**) a 24hr window of TTFields application. In the PRE-experiment, we found that the acute numbers of TMZ - TTFields-treated cells were similar to vehicle (Veh) -TTFields (i.e. group 4 vs group 3 in **Fig. 7E**). However the fold expansion of TMZ-TTFields treated cells seven days after treatment was significantly reduced compared to veh-TTFields (ie group 4 vs group 3 in **Fig. 7F**). In the POST-experiment, we also did not observe an acute reduction of TTFields-TMZ treated cells compared to control (i.e., group 4 vs group 3 in **Fig. 7H**), but did observe a significant reduction of TTFields-TMZ treated cells seven days after treatment (i.e. group 4 vs group 3 in **Fig 7I**). Surprisingly, in the ‘with’ experiment, we observed an acute increase in TMZ+TTFields-treated cells compared to veh+TTFields (group 4 vs group 3 in **Fig. 7K**), and a significant increase in the expansion of TMZ+TTFields co-treated cells 7 days after treatment (i.e. group 4 vs group 3 in **Fig. 7L**). Thus, TMZ given pre- or post-TTFields slows tumor cell recurrence, but co-administration enhanced tumor cell recurrence. We next asked if either of the treatment paradigms that slowed tumor cell recurrence correlated with the presence of ARL13B^+^ cilia. Since we observed a significant effect of giving TMZ after TTFields on parental cells, we examined fold expansions of S7 ARL13b KO cells 6 days after TTFields-TMZ treatment (**Fig. 7M,N**). However, there was no significant difference in the expansion between TTFields-TMZ-treated and TTFields-Veh treated ARL13B KO cells (i.e. group 4 vs group 3 in **Fig. 7N**). These results suggest the relative timing of TMZ exposure to TTFields impacts subsequent tumor cell expansion in vitro that also correlates with the presence of ARL13B^+^ cilia.

### 6. TTFields disrupt primary cilia in patient tumors ex vivo

While we currently do not have the technical capability to test TTFields in an intracranial, xenograft model, we asked if the effects of TTFields on cilia we observe in adherent or spheres of glioma cells are detectable within the patient tumor microenvironment. To test this, we divided fresh biopsy samples into 3 groups: immediate/acute fixation, 24hr control or 24hr of TTFields. We then fixed, and immunostained cryosections of biopsies (**Fig. 8A**). First we examined a subependymoma (a grade 1 glioma), a tumor type reported to possess cilia (49). We found that ARL13B^+^ cilia extending from OFD1^+^ basal bodies were readily detectable in control (**Fig. 8B**) whereas we only clearly observed OFD1^+^ basal bodies in the TTFields treated biopsy (**Fig. 8C**). We also received newly diagnosed GBM biopsies from a 34 year old male and 66 year old male which we treated similarly except immunostained basal bodies/centrioles with gTUB and cilia with ARL13B antibodies. We observed gTUB^+^ basal bodies with ARL13B^+^ cilia in acutely fixed tissue and in overnight control (**Fig. 8D, E, G**), whereas the cilia appeared blunted or generally reduced in TTFields treated tissue (**Fig. 8F, G**). Similar observations were made in a 66 year old male GBM biopsy exposed to 24hr of TTFields (**Suppl. Fig. 5**). These data are consistent with our cultured adherent cells and gliomaspheres, raising the possibility that TTFields disrupts ARL13B^+^ ciliated tumor cells within the tumor.

**Figure 8.**
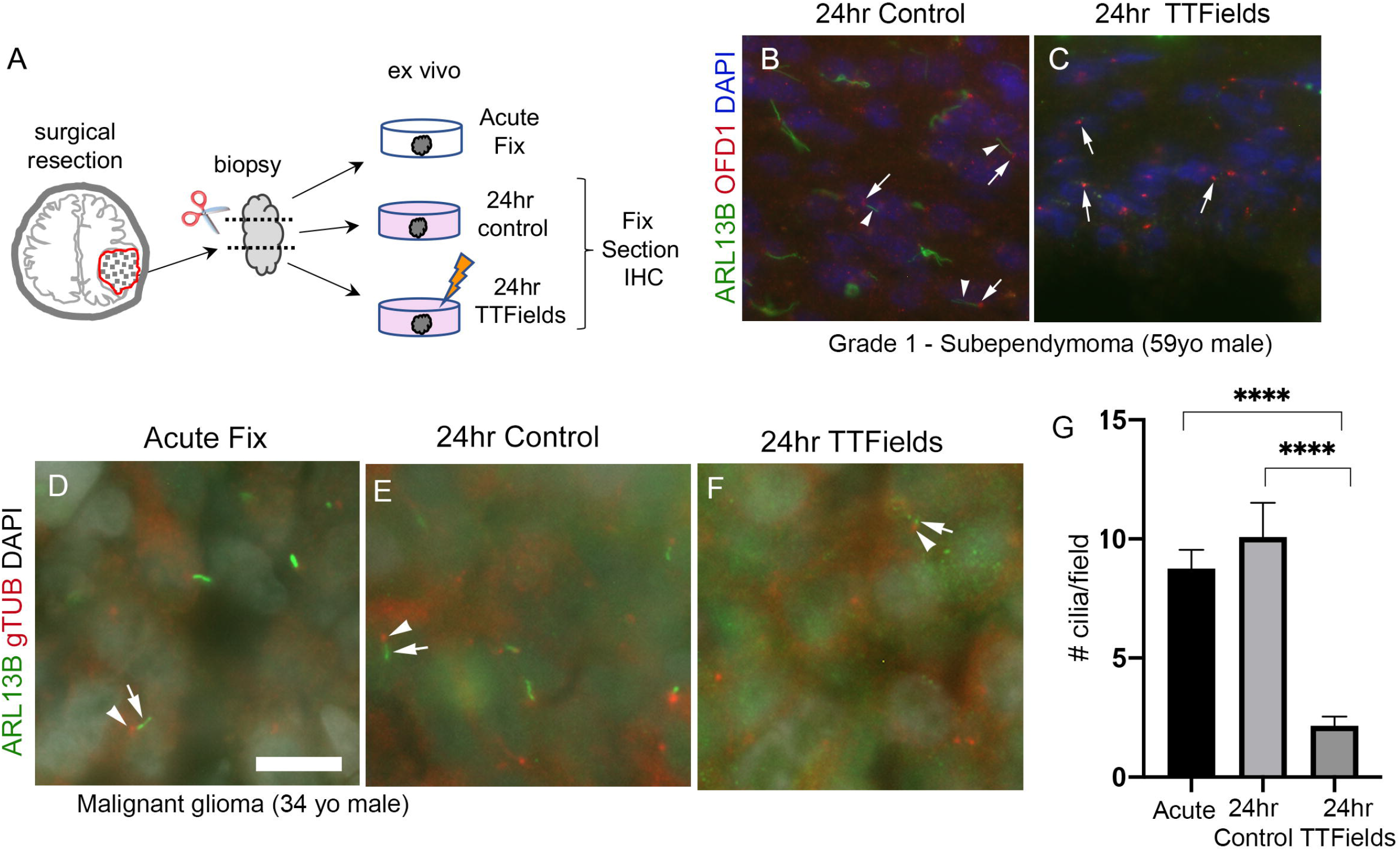
TTFields disrupt cilia in patient samples ex-vivo. **A**) Ex-vivo treatment of surgical resections. Biopsies were dissected and separated into immediate/acute fix, 24hr control or 24 TTFields treatment. Tissue was fixed, frozen, cryosectioned and immunostained. Nuclei are labeled with DAPI. **B**) Immunostaining of control section of a Grade 1 subependymoma showed ARL13B^+^ cilia (arrowheads) extending from OFD1^+^ basal bodies (arrows). **C**) TTFields-treated tissue section showed OFD1^+^ basal bodies (arrows) without obvious ARL13B^+^ cilia extensions. **D-F**) Tissue from a malignant glioma from a 34 year old male that was separated into immediate/acute fixation, 24 hour control, or 24 hour TTFields exposure. Tissues were fixed, cryosectioned and immunostained for ARL13B (green) and gTub (red), and nuclei labeled with DAPI (white). ARL13B^+^ cilia (arrows) with gTUB^+^ basal bodies (arrowheads) are readily detected in acute (**D**) and 24 hour control (**E**), but appeared blunted or generally absent in TTFields group (**F**). **G**) Mean (+/− SEM) number of cilia/field (n=12-13 fields/group) from samples in D-F. ****p<0.0001 (ANOVA). Scale bars (in μm) in **A** = 10.

## Discussion

We show that TTFields significantly impact the ability of glioma cells to maintain their primary cilium. Low and high grade glioma cells disassemble their cilia shortly after TTFields, though the population is not completely eliminated because ciliated cells reappear at similar frequencies if the treatment is stopped. The mechanism leading to cilia loss appears to involve a combination of autophagy activation and death of ciliated cells. TTFields does not similarly impact the cilia of various normal neural cell types, pointing to one aspect of glioma biology that may be differentially sensitive to TTFields. Further, TMZ-induced increase in ciliated glioma cells is inhibited by TTFields. This is of potential significance because ARL13B-mediated signaling associated with glioblastoma cilia is linked to both tumor growth and TMZ resistance in vivo (30). Thus disruption of cilia by TTFields may help enhance TMZ efficacy, or help eliminate a population of treatment-resistant cells. Surprisingly, we found that simultaneous TMZ and TTFields treatment is less effective on slowing tumor cell recurrence than TMZ added before or after TTFields, an effect that is lost on cilia-depleted tumor cells. Considering our observation that TTFields-mediated changes in cilia within patient tumors raises the possibility that tumors containing high levels of ciliogenesis could be more receptive to TTFields and TMZ, though future studies are needed to correlate tumor ciliation, TTFields and TMZ on patient outcome.

How does TTFields lead to glioma primary cilia dismantling? Within hours, TTFields trigger axonemal elongation with accompanying changes in the distribution of proteins along the ciliary membrane, culminating in the loss of the cilia. The re-distribution of ciliary membrane-associated proteins may be due to TTFields effect on membrane permeability reported to occur during the same timescale (16). It is not clear if TTFields cause cilia to be absorbed back into the cell, shed into the extracellular milieu, or both. Most mammalian cilia appear to disassemble by shedding the whole cilium (50, 51). However, live imaging studies of glioma cells during TTFields support a withdrawal or absorption back into the cell as we did not observe cilia detachment. Whatever the mechanism of cilia loss after TTFields, the changes in cilia may serve as a biomarker of TTFields efficacy in the tumor. Although TTFields eliminates cilia/ciliated glioma cells, they grow back at the same frequency though longer if treatment is stopped. We do not know if the regrown cilia are from the same cells, or represent a new population of ciliated cells that is more sensitive or resistant to treatment, which could potentially be addressed via extended live imaging during and after TTFields. The ciliary lengthening on recovered cells could be due to elevated autophagy pathway activation, which has been shown to elongate cilia (37). CQ has been observed to reduce the autophagy-mediated increase in cilia length on human kidney proximal tubular cells (52), which is further supported by our observation that CQ reduces the TTFields-induced ciliary lengthening (Fig. 4G, H).

Autophagy activation may represent one contributing factor in the disappearance of cilia after TTFields, since CQ pre-treatment did not fully restore the frequency of ciliated cells. TTFields may trigger many other factors that promote cilia removal. It is possible only a fraction of the cilia is removed by autophagy. HDAC6-mediated autophagy can result in ‘ciliophagy’ in mouse tracheal epithelial cells and cholangiocarcinoma cells (46, 53), and thus some glioma cilia may be driven by HDAC6 mediated autophagy whose signaling at cilia is a key regulator of glioma cell proliferation (32). Whatever the mechanism of autophagy linked to glioma cilia, it is unclear whether it is promoting or reducing cell survival. However, the link to autophagy is noteworthy as TTFields activation of autophagy appears to have dual significance. On one hand it may drive the death of cells (11), or alternatively promote activation of pathways that reduce sensitivity to TTFields (14). A scenario in which the ciliated glioma cells die by TTFIelds favors the former, whereas if glioma cells re-grow their cilia (supporting the return of cilia frequency) favor the latter. Alternatively, there could be a mixture of these scenarios that requires further study.

It is not clear why normal differentiating or proliferating neural cilia are less affected by TTFields. It is possible normal mouse neural cell type plasma membrane are more impervious to or recover quicker from the membrane permeating effects of TTFields than glioma ciliary membranes. For example, normal human fibroblast membranes were less perforated by TTFields than glioma cells (16). Neurons and glia however were not completely spared as cells showed lower frequencies with 3 days of exposure. However, the degrees of changes were far less than glioma cells. For example, neuronal cilia frequencies after 3 continuous days of TTFields were reduced by ~40% compared to control whereas the ciliary frequency in glioma cells was reduced by ~90%. The preservation of cilia from this stress could be cytoprotective. For example, primary cilia on neurons were recently reported to prevent neurite degeneration in developing cortical neurons after exposure to stressors in vivo including alcohol and ketamine (54). Similarly in normal glia, it was reported that hedgehog mediated signaling thru primary cilia promote cell survival in stressed in vitro conditions (55). Thus, the retention of cilia on neurons and glia may help protect against stress induced by TTFields. The extent to which TTFields affects tumor cells and normal neural cilia in the human brain will require post-mortem analyses.

Our findings raise the possibility that TTFields could help eliminate or suppress TMZ-resistant cell types. TMZ can stimulate expression of ARL13B and an interaction between ARL13B and the purine biosynthesis pathway as a mechanism to drive TMZ chemoresistance in glioblastoma (30). Thus, TMZ stimulation of ciliogenesis that we and others found, and increased sensitivity to TMZ in ARL13B KO cells, are consistent with the notion that cilia contribute to resistance to treatment. Thus, TTFields targeting of cilia or ciliated glioma cells may enhance TMZ toxicity. Given the opposing effects on TMZ and TTFields on ciliogenesis, we explored the effects of different order of these treatments and found that the interaction on subsequent proliferation depends on when TMZ is administered. Surprisingly we observed that simultaneous treatment of TTFields with TMZ worked against the efficacy of TTFields (Fig. 7L), through some unknown mechanism. However TMZ before or after TTFields application slowed tumor cell recurrence, an effect not observed in ARL13B-depleted cells. The implications may be two-fold. First the combined effects of TMZ and TTFields may depend on degree of ARL13B^+^ cilia in the tumor. Second, there may be a treatment window or boundaries surrounding TTFields, within which TMZ exposure may be counterproductive against slowing tumor recurrence. Whether TMZ and TTFields have converging actions at the cilium remains unclear. Both treatments appear capable of stimulating autophagy (14, 56, 57), yet their effects on ciliogenesis appear opposite. Nevertheless, autophagy pathway inhibitors during or subsequent to these treatments could help target cells that clearly survive both treatments.

Our study raises the possibility that gliomas with enhanced ciliogenesis potential may be more sensitive to TTFields therapy, a cellular susceptibility that may come with tradeoffs. A tumor containing more ciliated proliferating cells may be more impacted by TTFields than tumors with few ciliated cells. However, a complete or sustained ablation of cilia may generate cell offspring that are mutated or transform into other resistant cell types. Indeed, some GBMs and older glioma cell lines are or become cilia-devoid (26, 58, 59). In medulloblastoma, loss of cilia mediated due to ablation of *OFD1* can lead to SMO inhibitor treatment resistance and formation of ‘persister-like’ states that support tumor recurrence (60). While our study involves patient-derived cell lines and fresh patient surgical resections, a limitation of our studies is we are not yet able to examine TTFields effects on intracranial tumor growth in an animal model, a future direction of our studies. It is possible the changes in cilia may be less robust deep inside the brain. Nevertheless, the TTFields-mediated changes in cilia we observed ex vivo warrant these investigations at the organelle level. Because all available genomic or proteomic databases cannot inform about the degree of ciliation in a tumor, future studies will need to examine how survival of patient being treated with TTFields and TMZ therapy depends on the ciliated profile of their tumors.

## Acknowledgements

The authors would like to thank Drs. Y. Porat and M. Giladi at Novocure, Inc, and Drs. D Chen and D. Tran at University of Florida in the Dept of Neurosurgery for technical support and comments on the manuscript, and Dr. T. Caspary (Emory Univ.) for the pDest-Arl13b:GFP plasmid. Funding for this research was supported in part by a grant from the Florida Center for Brain Tumor Research and Accelerate Brain Cancer Cured (to M.R.S.), and a 2019 AACR-Novocure Tumor Treating Fields Research Grant (#19-60-62-SARK; to M.R.S.).

## Materials and Methods

### Cell culture

L0 (grade IV glioblastoma from a 43 year old male) and S7(grade II glioma from a 54 year old female with EGFR amplification) cell lines were isolated and maintained as previously described (26, 61, 62) (63). *ARL13B* and *KIF3A*-deficient L0 and S7 cells were generated using CRISPR/Cas9 as previously described (32). L0 and S7 cells were grown as floating spheres and maintained in NeuroCult NS-A Proliferation medium and 10% proliferation supplement (STEMCELL Technologies; Cat# 05750 and #05753), 1% penicillin–streptomycin (Thermofisher, Cat# 15140122), 20 ng/ml human epidermal growth factor (hEGF) (Cat #78006), and 10 ng/ml basic fibroblast growth factor (bFGF) (Cat #78003). For S7 cells, the media was supplemented with 2 μg/ml heparin (Cat #07980). All cells were grown in a humidified incubator at 37 °C with 5% CO2. When cells reached confluency, or spheres reached approximately 150 μm in diameter, they were enzymatically dissociated by digestion with Accumax (Innovative Cell Technologies; Cat#AM-105) for 10 min at 37 °C. For human cells grown on glass coverslips, NeuroCult NS-A Proliferation medium was supplemented with 10% heat inactivated fetal bovine serum (FBS) (Cytiva, Cat #SH30070.03HI).

Primary neural cultures were similar to previously described (32, 64). Briefly, acutely micro-dissected C57/BL6 mouse cortices from postnatal day 0-2 pups were dissected into Gey’s Balanced Salt Solution (Sigma-Aldrich, Cat #G9779) at ~37 °C under oxygenation for ~20 min. Dissociated cells were triturated with pipettes of decreasing bore size, pelleted by centrifugation at 1,500 rpm for 3-5 min, and resuspended and plated in glial medium containing DMEM (Cytiva HyClone, Cat# SH3002201), FBS (Gemini BioProducts, Cat# 50-753-2981), insulin (Sigma-Aldrich, Cat# 15500), Glutamax (Gibco, Cat# 35050061) and Penicillin-streptomycin (Gibco, Cat# 15140122). Cells were plated at a density of 80,000 cells/coverslip on 12-mm glass coverslips coated with 0.1 mg/ml poly-D-lysine followed by 5 μg/ml laminin in minimal essential medium. After approximately 2 hrs, cells were supplemented with 2mL neuronal media containing Neurobasal A (Gibco, Cat# 10888022) supplemented with B27 (Gibco, Cat# A3582801), Glutamax (Gibco, 35050061), Kynurenic acid (Sigma Aldrich, Cat# K3375), and GDNF (Sigma Aldrich, Cat# SRP3200). Every 4 days, half of the media was replaced with fresh neuronal media as described above but lacking kynurenic acid and GDNF. On DIV12, coverslips were transferred into TTFields dishes and fixed after 24 hrs or 3 days after treatment as described below.

### TTFields induction and Timelapse Imaging

For adherent and spheres, 5×10^4^ cells were seeded in 2ml growth media with or without 10% FBS, respectively. Adherent cells, spheres or biopsies were placed in TTFields ceramic dishes, each dish approximately the size of a single well of a 6-well plate, and mounted into inovitro™ base plates (Novocure Ltd., Haifa, Israel). The base plates were connected to a power generator which delivered TTFields at frequency of 200 kHz at a target intensity of 1.62V/cm (65). During TTFields treatment, cells were maintained in an incubator (ESCO Technologies, Horsham, PA) with the ambient temperature set to 18°C with 5% CO2 and a target temperature of 37°C inside each ceramic dish. Treatment duration are as indicated but ranged from 1 to 72 hours for a single treatment. To prevent media evaporation during TTFields application, parafilm was placed over each TTFields ceramic dish. In between repeated exposures or for recovery experiments, cells were dissociated and transferred back to a regular incubator. Control samples were grown at 37°C in 5% CO2 in 6 well plates. In some experiments, we pretreated cells before TTFields with either vehicle, or specified drugs. Unless otherwise stated, data in each experiment were pooled from at least 4 dishes per condition and per timepoint.

For timelase imaging combined with TTFields, we plated 50,000 cells in S7/L0 media supplemented with 5% FBS into 35 mm glass bottom culture dishes (Ibidi, cat #81158) which were maintained at at 37°C in 5% CO2. Twenty four hours before imaging at about 70% confluency, cells were transfected with 500 ng total cDNA/dish of pDest-Arl13b:GFP (gift from T. Caspary) and pCMV-myc/mCherry:hOFD1 (Vectorbuilder.com, vector ID: VB201119-1128fyp) using Lipofectamine 3000 (Life Technologies; Carlsbad, CA, cat#L3000015). A TTFields-delivering ceramic insert was placed into the culture dish and connected to a generator that delivered TTFields at a frequency of 200kHz at an approximate intensity of 1.2 V/cm and target temperature of 37°C inside each dish. Imaging was conducted on an inverted Zeiss AxioObserver D1 microscope using a Zeiss 40×/0.95 plan Apochromat air objective. The microscope stage was equipped with a Tokai Hit stage incubation system that maintained a humid environment and ambient temperature of 22-23°C and 5% CO2. Baseline images were captured every minute, whereas after TTFields onset, images were collected every 5 minutes with exposure times ranging in duration from 400 to 750msec (EGFP) and 300-400 msec (Cy3) per image. Image acquisition and processing were performed using Zeiss ZEN software.

### Ex-vivo culture and TTFields-treatment

In accordance with our institutional IRB protocol (#201902489), we collected several fresh, surgically-resected tumor biopsies that were subsequently pathologically confirmed. Within 1 hour of the resection, biopsies were taken to the laboratory, and dissected into several pieces using a sterile scalpel blade. Tissues were immediately fixed and/or transferred into 2mL of S7 media for culture at 37°C in 5% CO2 or transferred into TTFields dishes for a 24 hour exposure as described above. Following TTFields, control and treated samples were fixed and prepared as described above.

### Cell Growth, Viability Assays

For cell proliferation assay, cells (2.5-5 x10^4^) were seeded in 2 ml of growth media per well in 6-well plates, or in 1ml of growth media per well in 24 well plates for indicated duration. Cells were then treated with various drugs including chloroquine (CQ) (Sigma; Cat#C6628) (20μM diluted in sterile water), temozolomide (TMZ) (Sigma; Cat# T2577) (0.3 to 100 uM diluted in DMSO), 1,2-bis(o-aminophenoxy)ethane-N,N,N’,N’-tetraacetic acid (BAPTA) (Sigma; Cat# B1205) (1μM diluted in DMSO), or ethylene glycol-bis(β-aminoethyl ether)-N,N,N’,N’-tetraacetic acid (EGTA)(Sigma; Cat# RES3010E) (0.6mM diluted in DMSO). After indicated treatment durations, cells were enzymatically dissociated and replaced in 1X phosphate-buffered saline (PBS). Total cell counts were collected using a Bio-Rad TC20 automated cell counter. Bar graphs show the mean (+/− SEM) and were analyzed statistically using analysis of variance (ANOVA).

### Immunostaining

For immunocytochemical (ICC) and immunohistochemical (IHC) analyses, samples were fixed at indicated timepoints with 4% paraformaldehyde in 0.1 M phosphate buffer (4% PFA) for 30min or ice-cold methanol for 15minutes (ICC) to 1 hour (IHC) and washed with 1x PBS. Spheres or biopsies were cryoprotected in 30% sucrose in PBS followed by a 1:1: 30% sucrose and optimal cutting temperature compound (OCT) (Fisher Healthcare, #4585), frozen in OCT over liquid N2 and cryosection at 16μm. Samples were stained for the indicated primary antibodies (Table 1). Samples were incubated in blocking solution containing 5% normal donkey serum (NDS) (Jackson Immunoresearch; Cat#NC9624464) and 0.2% Triton-X 100 in 1x PBS for 1 hour and then incubated in primary antibodies with 2.5% NDS and 0.1% Triton-X 100 in 1x PBS either for 2 hours at room temperature (RT) or overnight at 4°C. For samples stained sequentially with mouse antibodies against gamma- and acetylated alpha-tubulin, samples were blocked with donkey anti-mouse IgG Fab fragments (20 μg/ml: Jackson Immunoresearch: Cat #715-007-003) as previously described (26). Appropriate FITC-, Cy3- or Cy5-conjugated secondary antibodies (1:1000; Jackson ImmunoResearch) in 2.5% NDS with 1x PBS were applied for 1-2 hour at RT, and coverslips were mounted onto Superfrost^TM^ Plus coated glass slides (Fisher Scientific, cat # 12-550-15) in Prolong Gold antifade media containing DAPI (Thermofisher; Cat# P36935). Stained coverslips were examined under epifluorescence using an inverted Zeiss AxioObserver D1 microscope using a Zeiss 40×/0.95 plan Apochromat air objective or a Zeiss 63X/1.4 plan Apochromat oil objective. Images were captured and analyzed using Zeiss ZEN software.

**Table 1.**
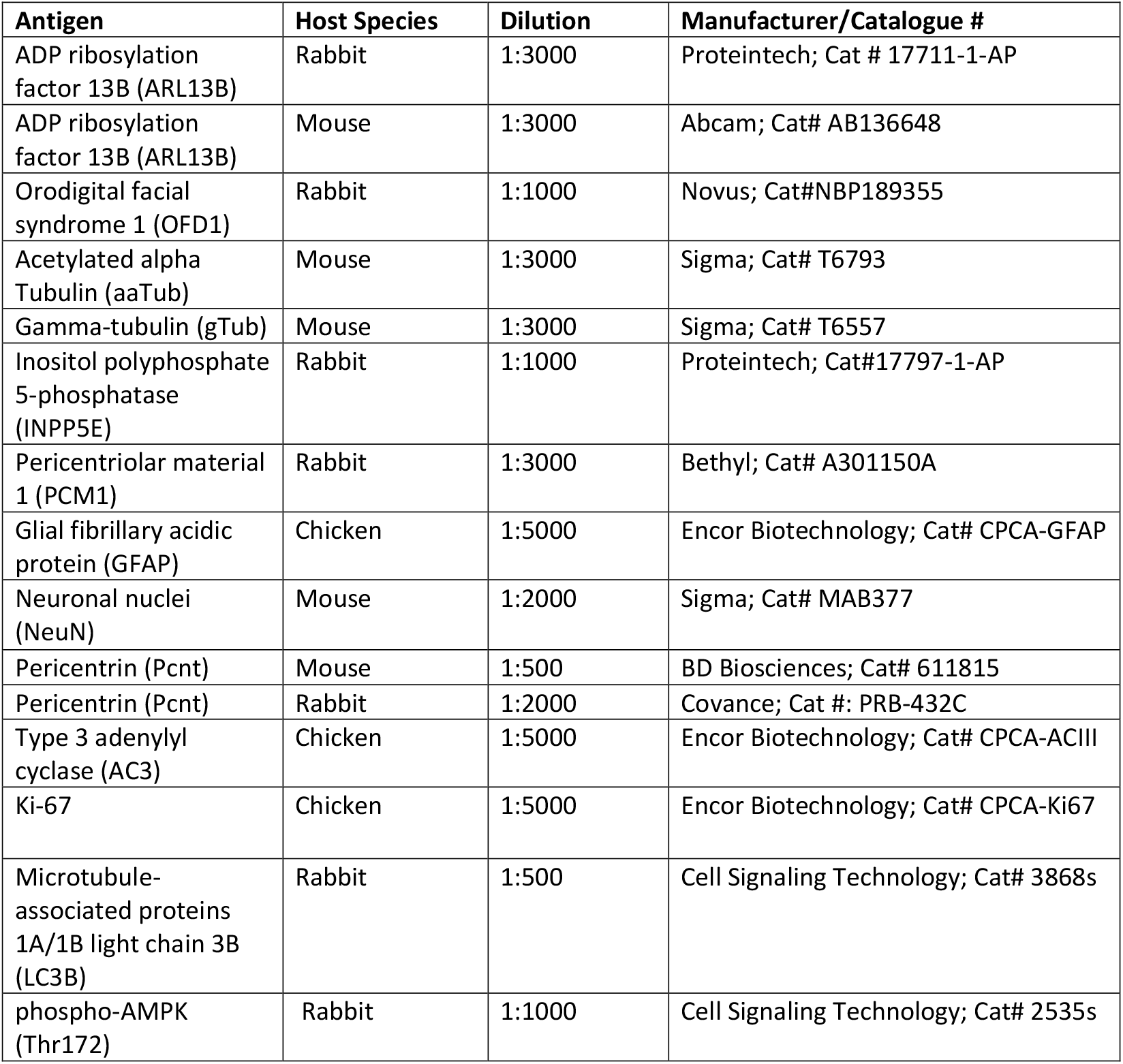
Primary antibodies used in this study.

