## Supplement Figures for "Tumor Treating Fields Suppression of Ciliogenesis Enhances Temozolomide Toxicity"

**Supplemental Figures and Legends**

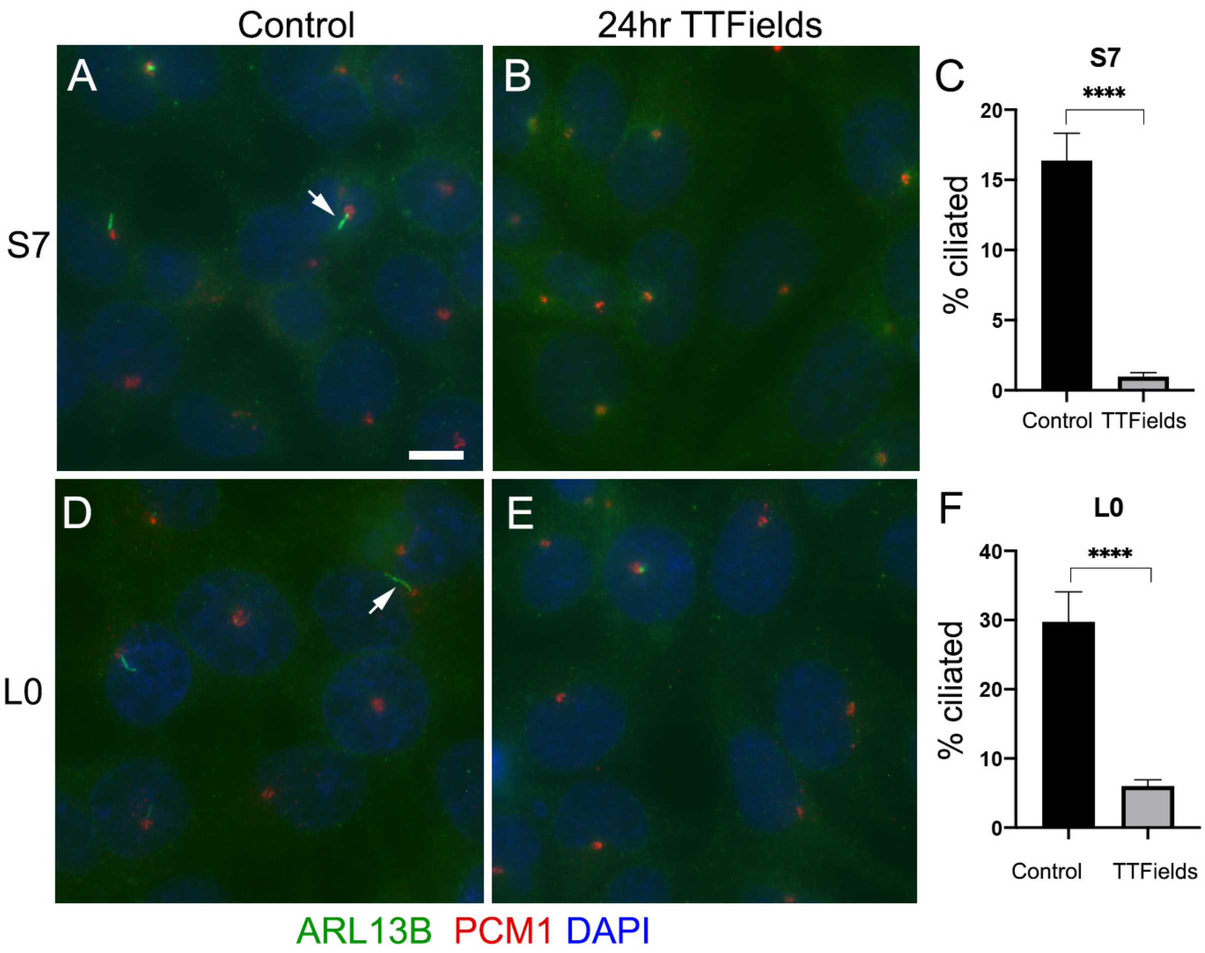


**Figure S1. TTFields ablates cilia within 24 hours in vitro**. **A,B**) S7 control (**A**) or TTFields-treated (**B**) cell stained for ARL13B (green), PCM1 (red). Nuclei are labeled with DAPI. **C**)Percentage of ARL13B⁺ cilia in S7 cells after 24 hours treatment. **D,E**) L0 control (**D**) or TTFields-treated (**E**) cell treated as in A,B. **F**) Percentage of ARL13B⁺ ciliated L0 cells after 24 hour treatment. ****p<0.0001 (ANOVA). Scale bars (in µm) in **A** = 10.

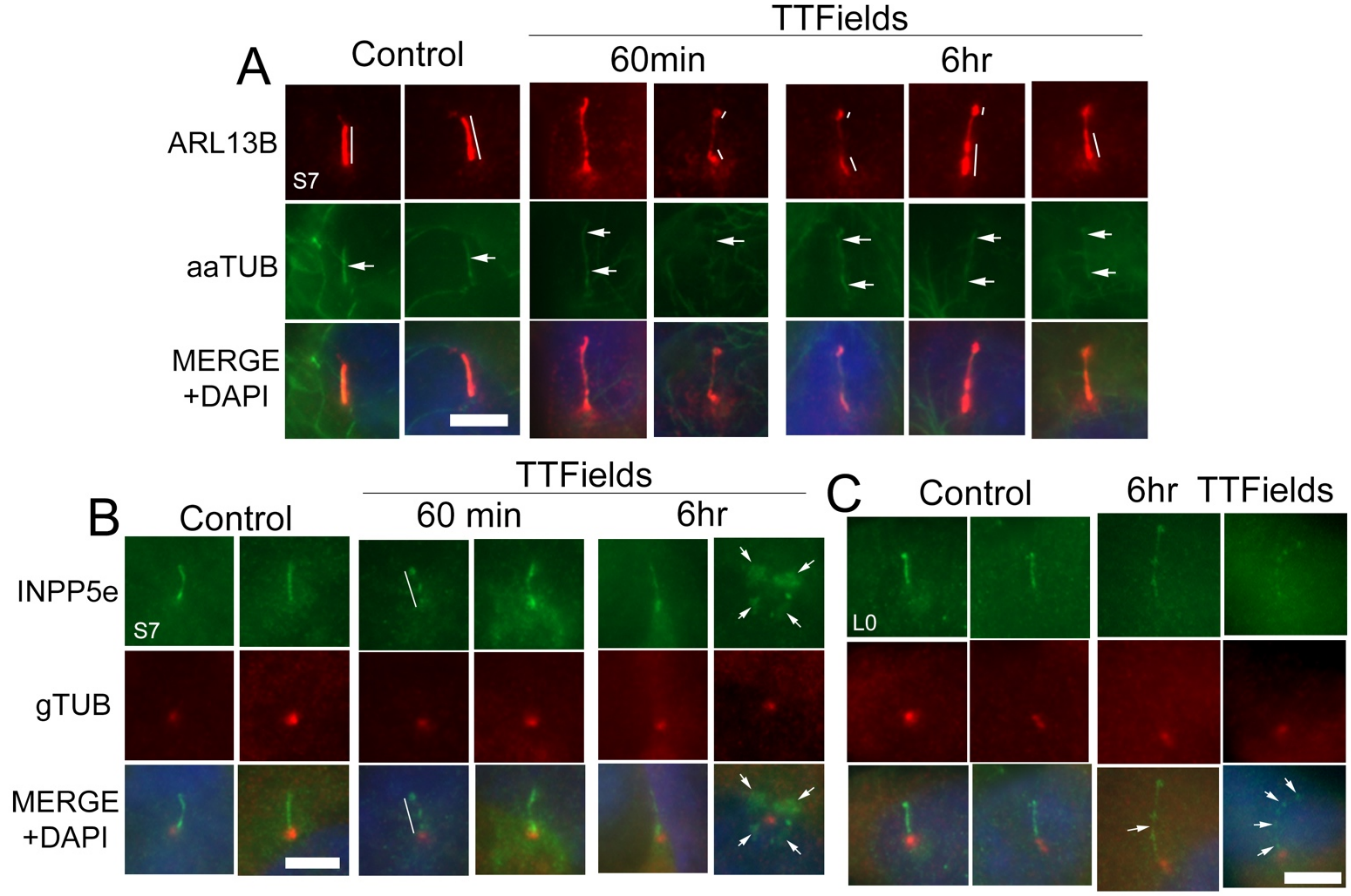


**Figure S2. Cilia elongation and alteration of ciliary membrane proteins by 6 hours of TTFields**. **A**) Adherent S7 control or TTFields-treated (60min or 6 hours) cells were fixed and immunostained for ARL13B (red) and aaTUB (green). In control, the distribution of ARL13B was relatively even along the length of the cilium (vertical lines). In TTFields exposed cells, the ARL13B distribution appears to polarize towards the base and distal tip. By 6 hours, TTFields cilia appeared longer with detectable underlying aaTUB$^{+}$ axonemes (arrows). **B**) Similar experiment as **A**, except we immunostained for INPP5e (green) which localizes to cilia membrane, and gamma-tubulin (gTUB, red) which is a component of the basal body/centriole. Like ARL13B, INPP5e distribution was largely evenly distributed along the length of the cilium but appeared puncta or polarized after TTFields. After 6 hours TTFields we also observed abnormal clusters of INNP5e (arrows) surrounding the gTUB$^{+}$ basal body/centriole. **C**) Adherent L0 control or TTFields treated cells stained for INPP5e and gTUB. TTFields-treated L0 cilia were also long and showed abnormal INNP5e distribution along the cilium. Scale bars in A, B and C = 5µm.

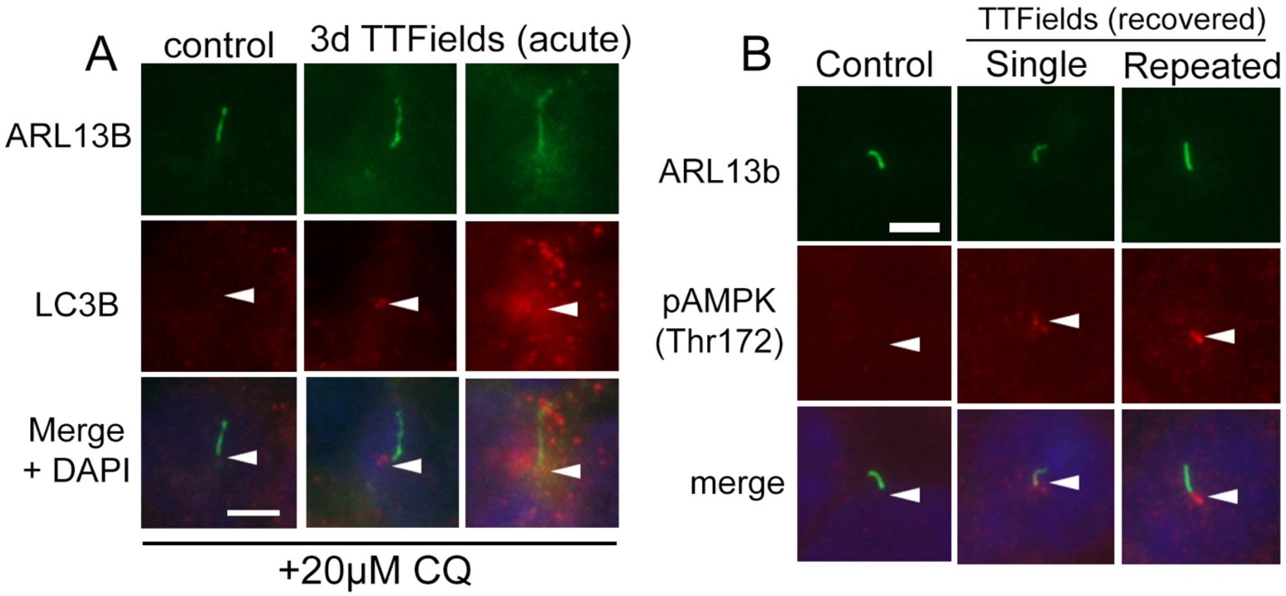


**Figure S3. Autophagy components LC3B and pAMPK localize to cilia after TTFields. A**) S7 control or TTFields-treated cells had 20µM chloroquine (CQ) added 3 hours before halting treatment and fixing cells. Cells were fixed and immunostained for ARL13B (green) and LC3B (red). TTFields treated cells showed recruitment or clustering of LC3B particles around the cilia/cilia base (arrowheads) compared to control. **B**) S7 control or TTFields-treated cells (either single 3 day continuous exposure) or repeated 3 days (as described in Figure 1A). Cells recovered for 4 days and were fixed and immunostained for pAMPK (Thr172), red). pAMPK signal was detected at the base of TTFields treated cilia compared to control (arrowheads). Scale bars in A and B= 5 µm.

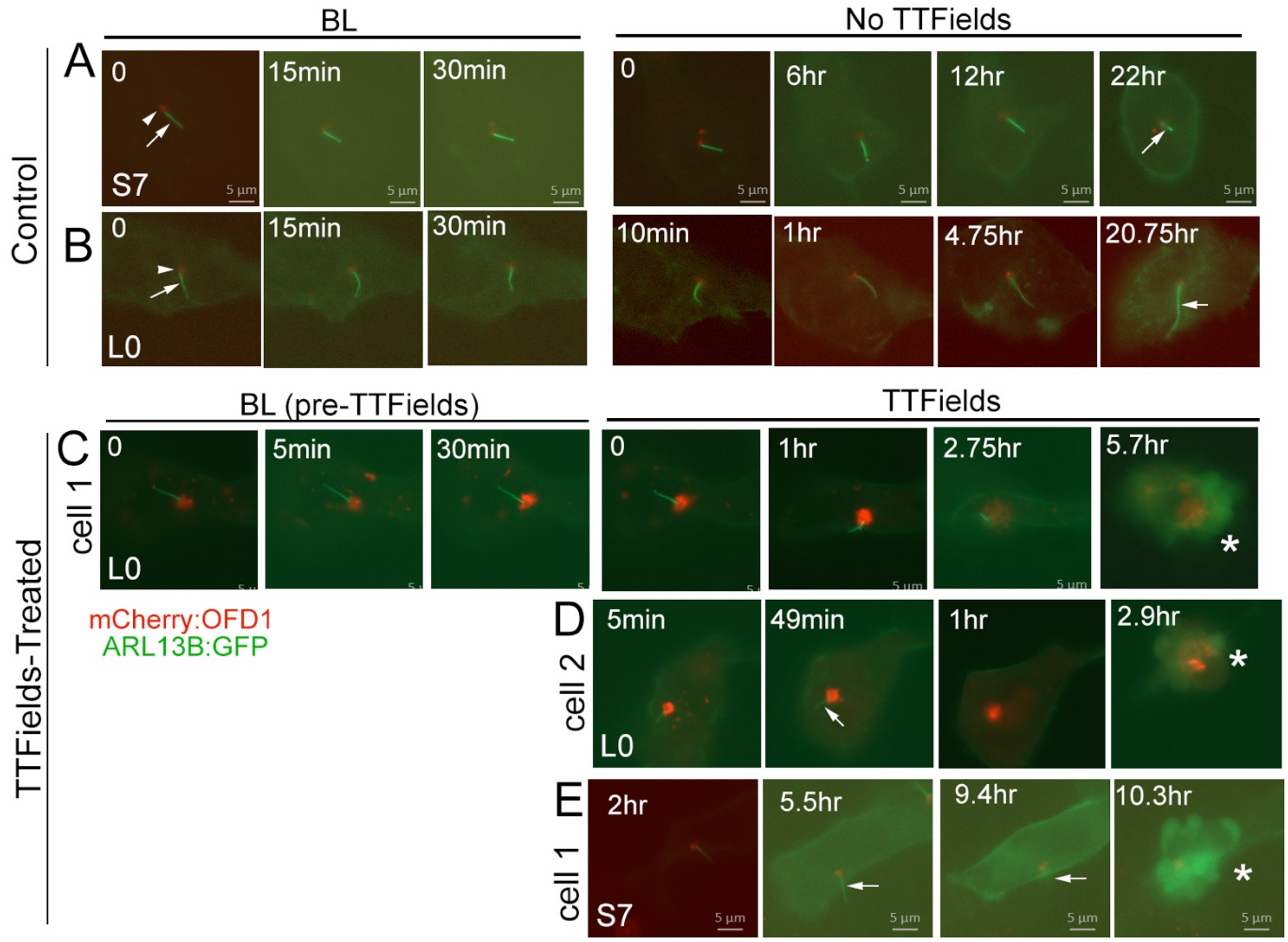


**Figure S4. Live imaging of glioma cilia during TTFields reveals death of ciliated cells. A,B**) Time-lapse imaging of S7 (A) and L0 (B) glioma cells transfected 24hr prior to imaging with cDNA encoding mCherry-tagged OFD1 (red) and GFP-tagged ARL13b (green) using lipofectamine. mCherry-OFD1 clusters around the basal body (arrowhead) whereas Arl13b:GFP enriches in the cilium (arrow). The row of images shows the cilium at indicated timepoints during baseline (BL) (images taken every minute) or overnight (images captured every 5 min) recording. **C**) Example of an L0 cilium during BL and TTFields. At 5.7hr the cell appears to die (asterisk). **D**) Another example of an L0 cell that appears to die during TTFields. **E**) Example of a ciliated S7 cell (e.g arrow at 5.5 hr) in which the cilium is not clearly visible at 9.4hr and the cell appears to die by 10.3 during TTFields. Bar = 5µm for all time lapse images.

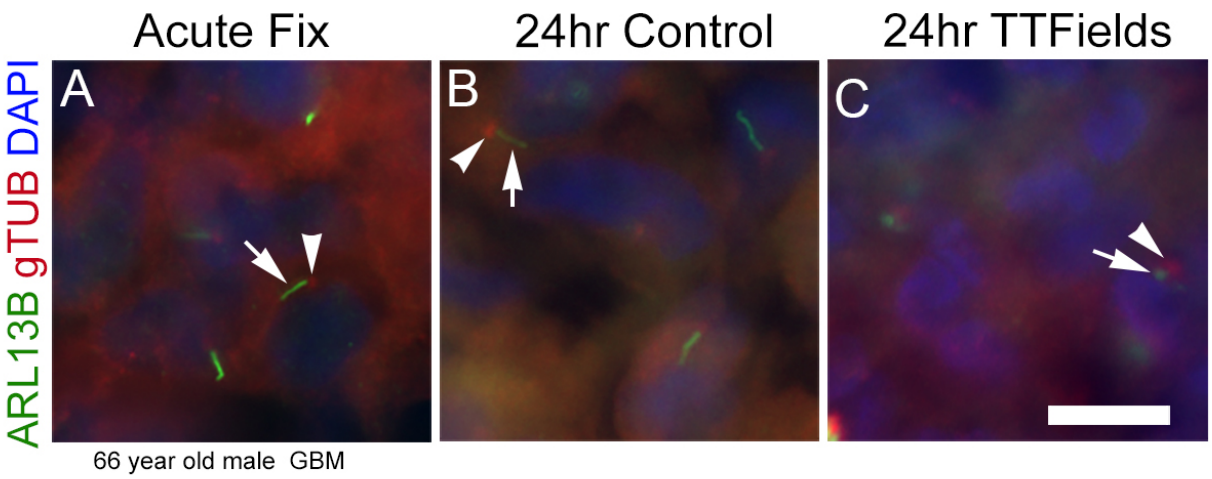


**Figure S5. Loss of cilia in a glioblastoma biopsy treated with TTFields ex vivo**. **A-C**) Fresh surgical resection from a glioblastoma from a 66 year old male was separated into immediate/acute fixation, 24 hour control, or 24 hour TTFields exposure. Tissues were fixed, cryosectioned and immunostained for ARL13B (green) and gTub (red), and nuclei labeled with DAPI (blue). ARL13B$^{+}$ cilia (arrows) with gTUB$^{+}$ basal bodies (arrowheads) are readily detected in acute (**A**) and 24 hour control (**B**), blunted or generally absent in TTFields group (**C**).
